# Metabolic Deficiencies Underlie Plasmacytoid Dendritic Cell Exhaustion After Viral Infection

**DOI:** 10.1101/2024.02.28.582551

**Authors:** Trever T. Greene, Yeara Jo, Monica Macal, Ziyan Fang, Fawziyah S. Khatri, Alicia L. Codrington, Katelynn R. Kazane, Carolina Chiale, Elizabeth Akbulut, Shobha Swaminathan, Yu Fujita, Patricia Fitzgerald-Bocarsly, Thekla Cordes, Christian Metallo, David A. Scott, Elina I. Zuniga

## Abstract

Type I Interferons (IFN-I) are central to host protection against viral infections^1^. While any cell can produce IFN-I, Plasmacytoid Dendritic Cells (pDCs) make greater quantities and more varieties of these cytokines than any other cell type^2^. However, following an initial burst of IFN-I, pDCs lose their exceptional IFN-I production capacity and become “exhausted”, a phenotype that associates with enhanced susceptibility to secondary infections^3–5^. Despite this apparent cost for the host, pDC exhaustion is conserved across multiple species and viral infections, but the underlying mechanisms and the potential evolutionary advantages are not well understood. Here we characterize pDC exhaustion and demonstrate that it is associated with a reduced capacity of pDCs to engage both oxidative and glycolytic metabolism. Mechanistically, we identify lactate dehydrogenase B (LDHB) as a novel positive regulator of pDC IFN-I production in mice and humans, show that LDHB deficiency is associated with suppressed IFN-I production, pDC metabolic capacity, and viral control following a viral infection, and demonstrate that preservation of LDHB expression is sufficient to partially restore exhausted pDC function *in vitro* and *in vivo*. Furthermore, restoring LDHB *in vivo* in exhausted pDCs increased IFNAR dependent infection-associated pathology. Therefore, our work identifies a novel and conserved mechanism for balancing immunity and pathology during viral infections, while also providing insight into the highly preserved but previously unexplained phenomenon of pDC exhaustion.

## Main

Persistent infections, including human immunodeficiency virus (HIV), hepatitis C virus (HCV), hepatitis B virus (HBV), *Mycobacterium tuberculosis*, and malaria represent massive burdens on human health. A key hallmark of these infections is immunosuppression, which limits responses against these pathogens, but also against unrelated secondary infections, cancer, and vaccination^6–8^. While immune suppression in this context can be beneficial to the pathogen, it also represents an adaptation by the host immune system that limits immunopathology and enables long-term host survival. Indeed, ablation of key immunosuppressive molecules often results in host death during infection even as control of the pathogen is improved^6,7^. Short-term alleviation of immunosuppressive adaptations has also shown therapeutic promise, allowing for clearance of pathogens and improved control of cancer^6,9–11^. These adaptations have been well studied in T cells and B cells where persistent stimulation promotes functional exhaustion^9,12^. However, relatively little is understood about the mechanisms that drive adaptations within the innate immune system.

Type I interferons (IFN-Is) are a family of key anti-viral and anti-neoplastic cytokines that act via limiting spread of viruses, restricting replication of transformed cells, as well as by enhancing innate and adaptive responses against viruses and tumors^1,13,14^. As specialized IFN-I producing cells, plasmacytoid dendritic cells (pDCs) rapidly respond to viral infection by synthesizing and releasing large quantities of IFN-I^2,15^. Through this function pDCs are known to promote the control of many viruses^2,15^. For instance, genetic deficiencies in virus sensing and IFN-I production by pDCs are associated with severe COVID-19 (reviewed in ^16^). However, after an initial robust response, pDCs lose their capacity to produce interferon in response to the ongoing infection or secondary stimulation^5,17^. This pDC phenotype, which is observed a few days after both acute and persistent viral infection^18^ and is sustained in the chronic setting, associates with compromised responses against unrelated secondary infections in both mice^3,4^ and humans^19,20^. Similar phenotypes have been described in several human and mouse cancer models as well ^21–24^ and promoting pDC IFN-I production capacity has been proposed as a means of overcoming cancer immunosuppression^25–28^. However, little is understood about the molecular processes that initiate and sustain pDC exhaustion in any model, and the evolutionary purpose of this highly conserved phenomenon is entirely unknown.

In this study, we investigate the transcriptional landscape of exhausted pDCs identifying a “pDC exhaustion signature” and changes in metabolic pathways that associate with their reduced IFN-I production capacity. By analyzing exhausted pDC metabolism, we determined that both oxidative phosphorylation (OxPhos) and glycolysis are suppressed in pDCs throughout a persistent viral infection. We uncovered that the metabolic enzyme lactate dehydrogenase B (LDHB) is downregulated in mouse and human exhausted pDCs, and ultimately demonstrated that LDHB supports IFN-I production in pDCs from both species. In line with this, chimeric mice with LDHB deficiency in the pDC compartment exhibited compromised viral control after coronavirus infection, demonstrating the importance of LDHB in pDCs for optimal antiviral responses. Mechanistically, we show that LDHB supports oxidative metabolism in pDCs, and is necessary to maintain normal ATP levels. Importantly, we demonstrated that restoration of LDHB in exhausted pDCs partially recovered their IFN-I production capacity *in vitro* and *in vivo* and that preserving pDC function in infected mice associates with IFNAR-dependent infection-induced pathology in the colon. Altogether, our results identify a key role for LDHB in the metabolism and function of pDCs and demonstrate that downregulation of LDHB and pDC exhaustion may have evolved to balance pDC antiviral function and immunopathogenic potential during infection.

### Transcriptional analysis of pDCs from virally infected mice identifies multiple pathways associated with their IFN-I exhaustion

We and others have previously described *in vivo* pDC exhaustion by using lymphocytic choriomeningitis virus (LCMV) infection in its natural murine host as a model system ^3,4,18^. While both acute and chronic infections with LCMV cause pDC exhaustion after a few days, only the persistent LCMV variant, Clone 13 (Cl13) sustains this phenotype for up to at least 30 days post infection (p.i.) ^3,4,18^. To characterize the transcriptional changes that take place in pDCs during infection, and highlight pathways associated with pDC exhaustion, we isolated pDCs from the spleens of uninfected mice or mice at 24 hrs, 8 days, or 30 days p.i. (Fig. 1a) and subjected them to RNA-seq. We observed many differentially expressed genes (DEGs) between uninfected and each timepoint (Adjusted p-value of less than 0.05, Log2 Fold Change greater than 1), with the largest number of DEG observed at 24 hrs p.i. (4248, Supplementary Table 1), fewer DEG at day 8 p.i. (2492, Supplementary Table 2), and the least number of DEG at day 30 p.i. (1787, Supplementary Table 3). We observed a high level of overlap in DEG identified for each timepoint with respect to uninfected pDCs ranging from 48% (when comparing day 1 and day 30) to 76% (when comparing day 8 to day 30) (Extended Data Fig. 1a). In line with this observation, principal component analysis (PCA) showed that, while each of these conditions clustered separately, day 8 and day 30 were more similar to each other than they were to day 1 p.i. (Fig. 1b). This likely represents differences between IFN-I producing pDCs (day 1 p.i.)^29^ and exhausted pDCs (day 8 and day 30 p.i.)^3,4,18^.

**Figure 1.**
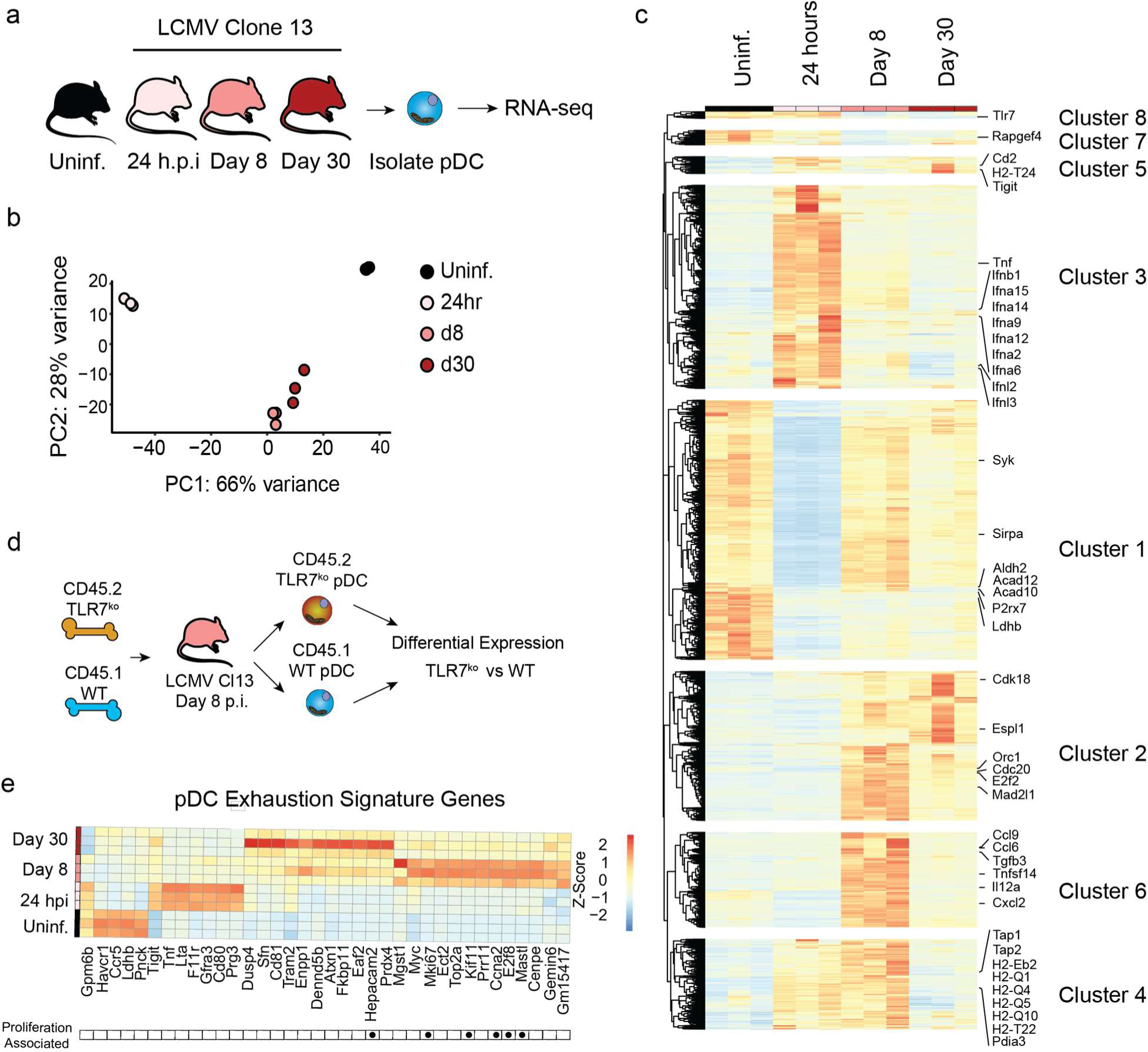
Defining the Transcriptome changes of pDCs throughout LCMV Infection. **(a)** Experimental design for pDC RNA seq. pDC (Lineage^-^, CD11c^+^, B220^+^, BST2^+^) were isolated from Uninfected or LCMV Cl13 infected mice at day 1 (24 hrs), 8, or day 30 p.i. and subjected to RNA-seq. **(b)**. Principal component (PC) plot showing pDC from uninfected mice, or from mice at day 1 (24 hrs), 8, or 30 p.i. **(c)** Hierarchical clustering of all genes that were DE between pDC from uninfected mice or from mice at day 1 (24 hrs), 8, or 30 p.i. **(d)** Experimental design for analysis of TLR7 dependent gene expression analysis. 50:50 TLR7^-/-^:WT mixed bone marrow chimeras were generated and infected with LCMV Cl13. At day 8 p.i., WT (Lineage^-^, CD11c^+^, B220^+^, BST2^+^, CD45.1^+^) or TLR7^-/-^ (Lineage^-^, CD11c^+^, B220^+^, BST2^+^, CD45.2^+^) were isolated and subjected to Microarray analysis. **(e)** Expression of genes differentially regulated in both TLR7^-/-^ vs WT pDC (from d) and in day 8 & 30-infected vs. uninfected mice (from a). Gene expression before and at different times after infection is depicted. Genes associated with cell cycle (GO: 0007049) are indicated by black dots.

Gene set enrichment analysis (GSEA) for DEG at 24 hrs. revealed 60 pathways significantly enriched (adj. p value<0.05) (Supplementary Table 4). These were largely associated with various viral and inflammatory diseases, pattern recognition, and cytokine production or signaling (e.g. Human papillomavirus infection, Human cytomegalovirus infection, Coronavirus disease, Epstein-Barr virus infection, HIV-1 infection, KSHV infection, Influenza A, Hepatitis B, Hepatitis C, Measles, Toll-like receptor signaling pathway, Cytokine-cytokine receptor interaction, RIG-I-like receptor signaling pathway) and are likely representative of the anti-viral response these pDCs are actively engaged with at this timepoint^3,29^. As expected, given their lower number of DEG pDCs from day 8 and day 30 showed fewer significantly enriched pathways when compared to pDCs from uninfected mice (18 and 10 respectively) (Supplementary Tables 5-6). At day 8 pathways were included several related to intracellular transport (e.g. regulation of actin cytoskeleton, axon guidance, synaptic vesicle cycle, nucleocytoplasmic transport) suggesting there may be changes in vesicle transport or the arrangement of the actin cytoskeleton of pDCs at this timepoint, and consistent with previously described morphological changes that pDCs undergo following stimulation^30^. We also observed several diverse metabolism related pathways (Fatty acid biosynthesis, Choline Metabolism in cancer, Cholesterol metabolism, Terpenoid backbone biosynthesis, glycosaminoglycan degradation) suggesting the metabolism of these cells may be altered. On the other hand, at day 30 p.i. we observed changes in metabolic pathways such as Vitamin B6 metabolism, Tyrosine metabolism, Linoleic acid metabolism, Taurine and hypotaurine metabolism, along with several disease related pathways that may represent the persistence of inflammation in this model (Legionellosis, Amyotrophic lateral sclerosis).

One strength of this data set is that it allows us to explore how pDC gene expression may change over time throughout infection. Therefore, to determine how changes in gene expression may regulate pDC function throughout infection we investigated the total set of DEG between the uninfected condition and all 3 infection timepoints (5361 unique genes). We then subjected these genes to hierarchical clustering analysis to identify temporally associated clusters of gene expression throughout infection (Fig. 1c). We then separated this into 8 clusters with unique temporal characteristics based on analysis of the within-cluster sum of squares as previously described^31,32^ (Extended Data Fig. 1b). For example, cluster 3 identified genes that were transiently increased at 24 hrs p.i. but returned to normal levels at day 8 and day 30. This includes most *Ifna* isoforms, *Ifnl2*, *Ifnl3*, *Ifnb1*, *Tnf*, and numerous other chemokines and cytokines (Supplementary Table 7) representing the acute anti-viral response that is subsequently suppressed in exhausted pDCs. Cluster 1 which represents genes that are mostly reduced through the entire course of infection (Extended Data Fig. 1b) includes many metabolism related genes (e.g. Aldh2, Acad12, Acad10, P2rx7, Ldhb and others, Supplementary Table 7). Cluster 2 which represents genes increased at days 8 & 30 p.i. is highly enriched for genes associated with cell-replication (e.g. Orc, Cdk18, Cdc20), in line with our previous observations that pDCs at day 8 and day 30 gain proliferative potential^18^. Cluster 4, which is made up of genes that are increased in expression at 24 hrs p.i. and 8 days p.i., but not at day 30 p.i. includes many antigen presentation associated proteins (Tap1, Tap2, H2-Eb2, H2-T22), in line with previous observations that pDCs gain some antigen presenting potential following initial stimulation^2^. Finally, cluster 6 includes genes specifically high at day 8, but not other timepoints. This notably contains a set of cytokines and chemokines which are not highly upregulated during the acute response at 24 hrs p.i., but are at day 8 (e.g. Tgfb3, Il12a, Ccl6, Ccl9, Cxcl2).

We then sought to identify genes and pathways that associate specifically with pDC exhaustion. For this we investigated the overlap between DEG at day 8 p.i. and day 30 p.i. as compared to pDCs from uninfected mice. We subjected this gene list to overrepresentation analysis of gene ontology (GO) terms. To restrict to the most significant pathways we considered only pathways with 5 or more genes identified, and an adjusted p-value less than 0.1. This identified 1260 pathways (Supplementary Table 8), though notably these were largely overlapping in scope with many, as expected based on previous work, being related to cellular proliferation, and anti-pathogen responses. To refine this analysis, we then made use of simplifyEnrichment^33^ to perform semantic similarity analysis and identify highly similar pathways. Using the binary cut method^33^ to determine clusters, this analysis identified 16 clusters of GO pathways (Supplementary Table 8) which share semantic similarity. Of these, 13 were large enough to be associated with common keywords shared by those pathways (Extended Data Fig. 1c), while the remainder included too few pathways for this type of analysis. As expected, several clusters were enriched for terms relating to the cell cycle and survival (e.g., chromosome, segregation, cycle, mitotic, checkpoint, proliferation, apoptosis), as well as a cluster enriched for terms relating to immune responses (immune, lymphocyte, interleukin). There were also several clusters related to cell-cell interaction (adhesion, cell-cell, junction, integrin), potentially relating to the role of pDCs in forming interferogenic-synapses during infection^34–42^. Finally, we also found a collection of pathways relating to cellular metabolism (metabolic), again reinforcing the potential for changes in metabolism in exhausted pDCs. Altogether these data bolster previous observations that pDCs gain proliferative capacity^18^, and increase apoptosis^43^ following infection. Furthermore, these data suggest that exhausted pDCs may have alterations in their cell-cell interactive properties and/or metabolic capacities which have not yet been described.

We then sought to further refine our analysis to specifically increase the proportion of pDC exhaustion associated genes. We previously demonstrated that pDCs from LCMV infected mice became functionally exhausted, in part, through persistent and cell-intrinsic signaling via TLR7^18^. Thus, to identify pDC exhaustion related genes we analyzed the differences in pDC gene expression in TLR7 knockout (TLR7^-/-^) and wild type (WT) pDCs from LCMV Cl13 infected mixed bone marrow (BM) chimeras (Fig. 1d), in which both WT and TLR7^-/-^ pDCs are exposed to the same infectious environment^18^. In this case the WT pDCs are exhausted, while the TLR7^-/-^ pDCs retain their function^18^. Direct analysis between these groups identified 110 DE genes, (Adjusted p-value <0.1, Log2 (Fold Change)) with magnitude more than 0.75, Supplementary Table 9).

We then compared these genes to the set of genes that are persistently differentially expressed between pDCs from uninfected (No Inf.) animals (functional) and those from day 8 or day 30 (exhausted). Altogether this identified 36 genes that met these criteria. This “pDC exhaustion signature”, includes several notable genes that have previously been observed to be altered in exhausted pDCs during SIV infection (*Tigit, Ccr5*)^44^, as well as molecules known to regulate pDC function (*Cd81*^45,46^, *Myc*^47^) (Fig. 1e). Additionally, this signature contains several cell-cycle related genes (GO: 0007049, Fig. 1e), in line with our previous observation that acquired proliferative capacity in pDCs is partially TLR7 dependent^18^. All together these data provide a resource of gene expression data for pDC exhaustion, including the first defined pDC exhaustion signature which may be useful for the identification of pDC exhaustion related phenotypes in future studies.

### LCMV infection drives a persistent reduction in metabolic capacity of pDCs

It has previously been appreciated that metabolism regulates pDC capacity to produce IFN- I^48–50^, however no study has so far investigated the long-term impact of viral infection on pDC metabolism. Given our analysis identified metabolism as a recurrent theme in pathways changed in exhausted pDCs (Supplementary Tables 5,6, Extended Data Fig. 1c), we sought to determine if metabolism of exhausted pDCs was significantly altered during infection. For this we used the Agilent Seahorse platform to evaluate oxygen consumption rate (OCR) as a proxy for OxPhos, and extracellular acidification rate (ECAR) as a proxy for lactate terminal glycolysis^51^. C57BL/6 mice were infected with LCMV Cl13 or left uninfected (Uninf.). On day 9 or 30 (p.i.), we isolated pDCs from the BM of these mice and evaluated their basal OCR and ECAR. We observed a significant reduction in both basal ECAR (Fig. 2a,c) and basal OCR (Fig. 2b,d) in pDCs from infected mice in both the acute (day 9 p.i.) and chronic (day 30 p.i.) phases of LCMV Cl13 infection. This was coordinate with a reduction in ATP production defined as the amount of OCR lost after ablating ATP synthase activity with Oligomycin (Fig. 2e). Spare respiratory capacity (SRC) or the difference between basal OCR and the maximal OCR measured by decoupling mitochondrial electron transport with FCCP (Fig. 2f) was also reduced. This suggests both baseline OxPhos activity and pDC potential for oxygen consumption are severely restricted in exhausted pDCs compared to their functional counterparts. Intriguingly, we did not observe any differences in OCR in the first 3 hrs following stimulation of either functional or exhausted pDCs with CpG-A (Extended Data Fig. 2a), this differs from what has been observed in humans which have showed an increase in OCR following stimulation with either HSV or Influenza Virus^50^ and may represent either a functional difference between mouse and human pDCs or a difference in response to diverse stimuli. As a parallel measure of mitochondrial activity we assessed levels of mitochondrial associated superoxide (mtSOX) in pDCs using a superoxide specific mitochondrially associated dye (MitoSOX™ Red), and found that mtSOX was much lower in pDCs from LCMV Cl13 infected mice at both day 9 and day 30 p.i. (Fig. 2g,h). Thus, our results suggest that the capacity to engage OxPhos as well as glycolysis is compromised in exhausted pDCs.

**Figure 2.**
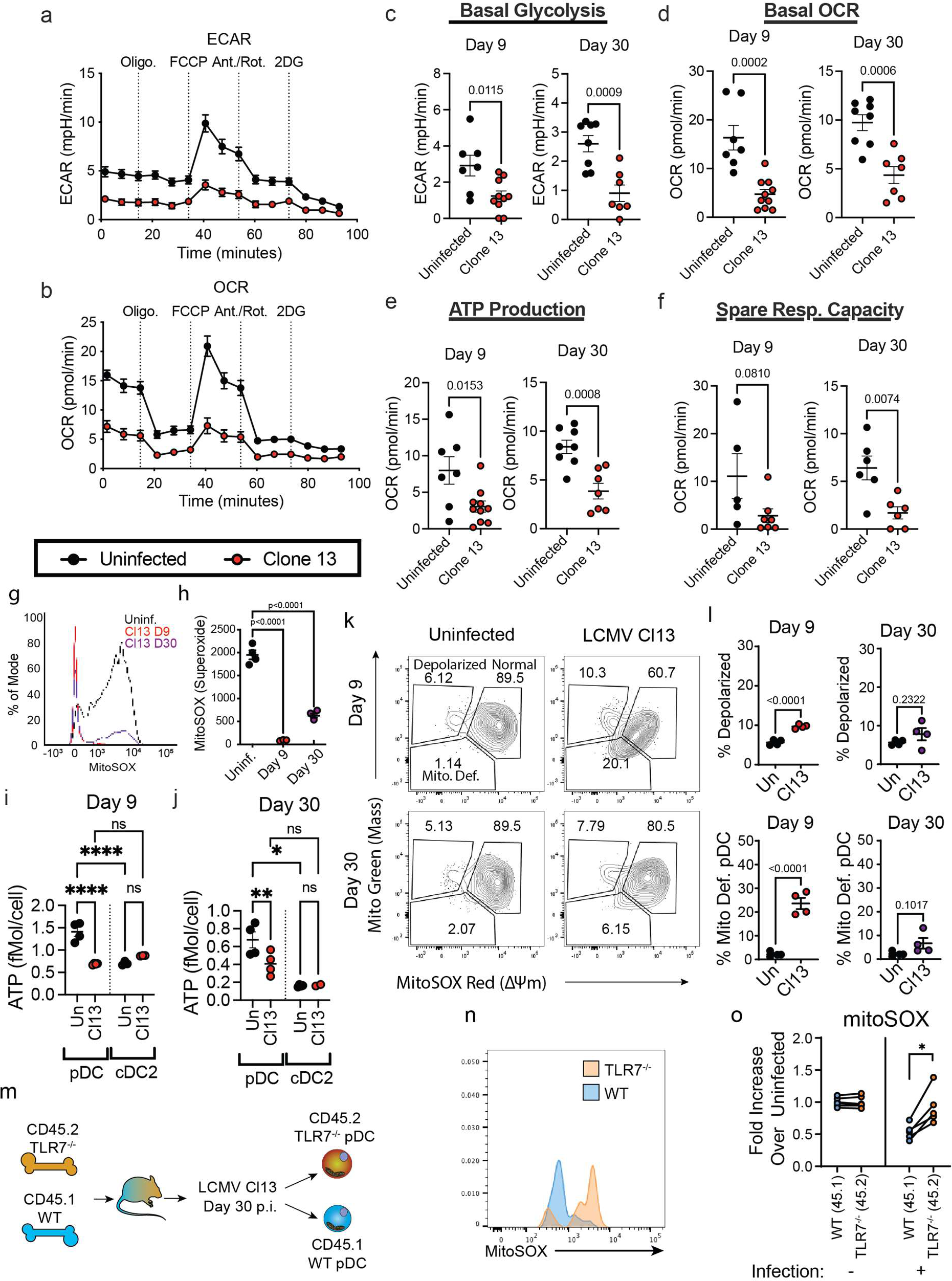
Clone 13 infection drives long-term changes in metabolism of pDCs and short-term changes in mitochondrial content. **(a,b)** Seahorse assay traces of extracellular acidification rate (ECAR)**(a)** and oxygen consumption rate (OCR)**(b)** of pDCs isolated from uninfected (black) or LCMV Cl13 infected (red) mice at day 30 p.i. **(c-f)** Derivative measures of ECAR and OCR from Seahorse assay at day 9 and day 30 p.i. (red) compared to uninfected mice (black) **(c)** Basal Glycolysis **(d)** Basal OCR **(e)** ATP production, **(f)** Spare Respiratory Capacity **(g)** representative stain of MitoSOX in pDC from uninfected mice (black) and LCMV Cl13 infected mice at day 9 (red) or day 30 p.i. (purple) **(h)** Quantification of MitoSOX gMFI from pDCs. **(i,j)** Total per-cell ATP content in pDC and cDC2 from uninfected mice or LCMV Cl13 infected mice at day 9 **(i)** or day 30 p.i. **(j)**. **(k)** Representative staining of mitochondrial mass (MitoGreen) and charge (MitoSox) from pDCs at indicated days p.i. **(l)** Quantification of percentage of pDCs with depolarized mitochondria (top) or that are mitochondria deficient (bottom) at indicated days p.i. **(m)** Experimental outline for panels n,o. 50:50 TLR7^-/-^:WT mixed bone marrow chimeras were generated and infected with LCMV Cl13. At day 30 p.i., WT (Lineage^-^, CD11c^+^, B220^+^, BST2^+^, CD45.1^+^) or TLR7^-/-^ (Lineage^-^, CD11c^+^, B220^+^, BST2^+^, CD45.2^+^) pDCs were analyzed for MitoSox staining via FACS. **(n)** Representative staining of MitoSOX in TLR7^-/-^ and WT pDCs from m, **(o)** Quantification of MitoSOX levels in WT & TLR7^-/-^ pDCs from m. Data are pooled from 3-5 experiments **(a-f,i,j,o)**, or representative of two independent experiments **(g,h,k,l)**. Data are shown as mean ± SEM. Statistics used are Student’s T Test **(c-f,l)**, paired T test **(o),** One way ANOVA with Tukey Correction (**h**), or Two way ANOVA with Fisher’s LSD test. p<0.05 *, p<0.01 **, p<0.001, ***, p<0.0001,****.

We then sought to determine whether ATP levels were altered in exhausted pDCs. For this we used a luminescence-based assay of ATP content, comparing pDCs from uninfected mice with pDCs from infected mice at day 9 or day 30 p.i. (Fig. 2i,j). We observed a striking decrease in total ATP content in pDCs at both days in infection, supporting our Seahorse assay observations and further demonstrating that exhausted pDC are metabolically unfit. Importantly, this is not a general phenomenon as conventional (c)DC from these same mice did not show any appreciable difference in ATP content (Fig. 2i,j). Notably, we also observed that, on a per-cell basis naïve, but not exhausted, pDCs showed higher levels of ATP than cDCs (Fig. 2i,j).

To determine if the changes in metabolism we observed in pDCs during chronic viral infection were the result of changes in mitochondrial content or function we assessed mitochondrial content (mtMass) and mitochondrial charge (mtΔψ) by flow cytometry. Mitochondrial mass and charge were evaluated together using charge dependent (MitoTracker™ Red chloromethyl-X-rosamine (CMXRos)) and independent (MitoTracker™ Green FM) dyes. In uninfected animals the majority of pDCs had similar mtMass and mtΔψ, with a small population showing high mass and low mtΔψ likely representing pDCs with depolarized mitochondria (Fig. 2k, top-left), as has previously been described in exhausted T cells ^52^. This population expanded significantly in the early stages (day 9 p.i.) of LCMV Cl13 infection (Fig. 2 k,l), but was not significantly different at day 30 p.i (Fig. 2 k,l), although average levels were still slightly higher than baseline at this later time point. Additionally, during LCMV Cl13 infection a third population emerged, showing low mtMass and mtΔψ as compared to pDCs from uninfected mice (Fig. 2k, top-right). As this population retained proportionality between mtMass and mtΔψ it is unlikely these are depolarized, but instead have reduced mitochondrial mass (Fig. 2 k,l). We refer to these cells as mitochondria deficient pDCs. By day 30 p.i. the number of mitochondria deficient pDCs were again low, although this remained slightly but not significantly higher than baseline (Fig. 2l). Therefore, changes in mitochondrial content may at least partially explain the loss of OxPhos observed at day 9 after LCMV Cl13 infection but are unlikely to fully explain the sustained loss of OxPhos that associates with long-term pDC exhaustion observed at day 30 p.i.

While these data correlate loss of OCR and ECAR in mitochondria to the timeline of exhaustion, we sought to determine if these changes were also dependent on TLR7. For this we again generated TLR7^-/-^:WT mixed BM chimeras, and analyzed the level of mtSOX in each compartment at day 30 p.i. (Fig. 2m). We observed no differences in mtSOX levels in WT and TLR7^-/-^ pDCs prior to infection, but mice infected with LCMV Cl13 showed reduced levels of mtSOX specifically in the WT compartment (Fig. 2n,o) suggesting that, like functional exhaustion ^18^, loss of mtSOX production is cell intrinsically dependent on TLR7 signaling in pDCs.

### LDHB in pDCs promotes IFN-I production and Viral Control

Given the reductions in pDC OxPhos and glycolysis during sustained infection we sought to investigate genes within the aforementioned “pDC exhaustion signature” (Fig. 1e) that have established roles in glycolysis as well as the TCA cycle and/or electron transport (Reactome R-MMU-1428517.1). Among the genes in our pDC exhaustion signature (Fig. 1e) only Lactate dehydrogenase B (*Ldhb*) met these criteria. Additionally, *Ldhb* is specifically highly expressed in pDCs as compared to cDCs in uninfected mice (Fig. 3a), and it is strongly and persistently downregulated during LCMV infection (Fig. 3a, Extended Data Fig. 2b) suggesting it may represent a reasonable candidate regulator of pDC function. LDH enzymes composed of LDHB show slower Vmax for pyruvate reduction to lactate than those comprised primarily of LDHA, and a strong substrate inhibition by pyruvate^53,54^. As such LDHB is primarily expressed in aerobic tissues where it is thought to promote OxPhos *via* the conversion of lactate to pyruvate^53^, although it is important to also acknowledge that under specific conditions LDHB also supports lactate terminal glycolysis^55,56^. Notably, we observed no significant change in the expression of *Ldha* in pDCs throughout the course of infection (Supplementary Tables 1-3).

**Figure 3.**
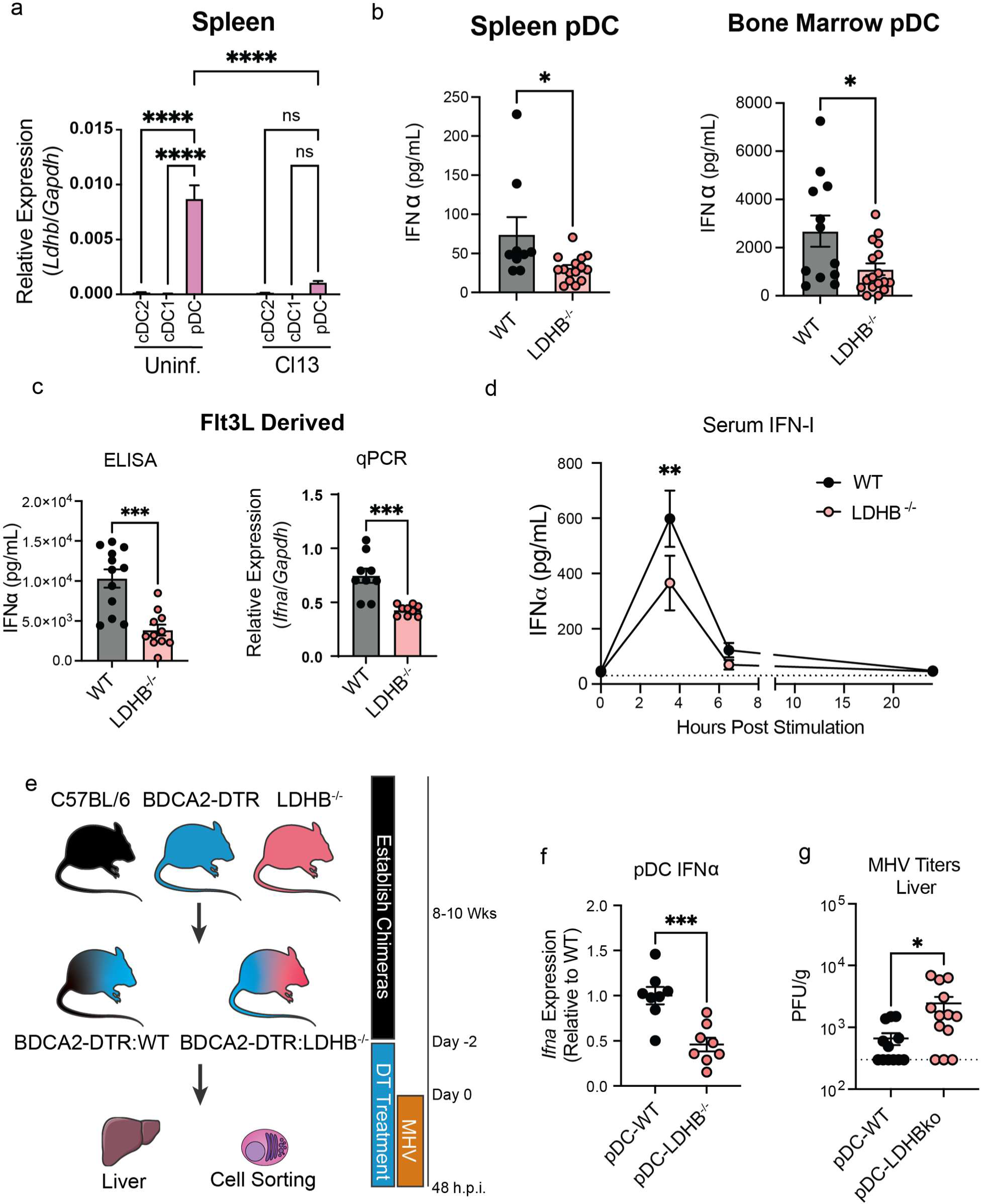
LDHB is essential for optimal mouse pDC IFN-I production *in vitro* and *in vivo*. **(a)** Relative expression of LDHB was evaluated by RT-qPCR in purified cDC2, cDC1, and pDC from the spleens of uninfected and LCMV Cl13 infected mice at day 9 p.i. **(b)** ELISA quantification of IFNα secreted by pDC isolated from the spleen (left) or bone marrow (right) of WT or LDHB^-/-^ mice after 12hr stimulation with CpG-A. **(c)** ELISA (left) or qPCR (right) quantification of secreted IFNα or *Ifna* transcript in pDC isolated from Flt3L cultures derived from WT or LDHB^-/-^ mice after 12hr stimulation with CpG-A. **(d)** ELISA quantification of IFNα in the serum of WT (black) or LDHB^-/-^ (red) mice treated with CpG-A. **(b-d)** Unstimulated pDC were below the limit of detection. **(e)** Experimental design for f-g, 50:50 mixed bone marrow chimeras for BDCA2-DTR:WT or BDCA2-DTR:LDHB were generated, then treated with DT daily from two days prior to infection to deplete BDCA2 expressing pDCs. Mice were then infected with MHV, 48 h.p.i pDCs were sorted and virus titers were quantified. **(f)** Relative expression of *Ifna* transcript in pDC from e. **(g)** Plaque assay quantification of MHV in livers from e. Data are pooled from 2-4 **(b-d,f,g)** independent experiments or representative of 2 independent experiments **(a)**. Data are shown as mean ± SEM. Statistics used are One way ANOVA with Tukey Correction **(a)**, Student’s T test **(b-d,f,g)**. p<0.05 *, p<0.01 **, p<0.001 ***,p<0.0001 ****.

Given that there is no relationship defined for LDHB with respect to pDC function we sought to determine if LDHB regulated pDC IFN-I production. For this we used LDHB knockout mice (LDHB^-/-^) generated by the International Mouse Phenotyping Consortium^57^. We first characterized the DC compartments of these mice and found no differences in total pDC or cDC numbers or their MHC-II expression, indicating that LDHB deficiency does not alter the number or maturation of DCs at steady state in the spleen (Extended Data Fig. 2c,d) or BM (Extended Data. Fig. 2e,f).

To then investigate whether DC function was impacted by LDHB deficiency we FACS-purified pDCs or cDC2s from the spleens of LDHB^-/-^ animals or age matched WT controls, stimulated them with CpG-A and measured IFNα by ELISA. Under these conditions LDHB^-/-^ pDCs from the spleen and BM produced less IFNα than their WT counterparts (Fig. 3b), despite no changes in cell viability (Extended Data Fig. 2g). We observed similar results when pDCs were derived from day 8-Flt3L-cultures of LDHB^-/-^ BM (Fig. 3c), demonstrating that the aforementioned IFN-I reduction was not due to a potentially different metabolic environment in LDHB^-/-^ mice. Importantly, conventional (c)DCs isolated from the same animals or from Flt3L- cultures showed no change in their IFNα production capacity (Extended Data Fig. 2h,i), demonstrating a pDC specific requirement for LDHB in IFN-I production. Additionally, quantification of *Tnfa* transcript in Flt3L derived pDCs showed no reduction in production of this cytokine (Extended Data Fig. 2j). This suggests there may be differences between LDHB deficiency and the state of pDC exhaustion, as exhausted pDCs can also show a defect in their capacity to produce TNFα ^18^.

We then moved to determine if this observation could be extended *in vivo*. We made use of the synthetic TLR9 ligand CpG. IFN-I responses to CpG in mice peak quickly at 3.5 hrs and are pDC dependent ^58,59^. WT or LDHB^-/-^ mice were injected with CpG, and IFN-I levels monitored in the serum. Despite similar number of pDCs in spleens and BM of WT and LDHB^-/-^ mice (Extended Data Fig. 2c,e), we observed significantly lower quantities of IFN-I in the serum of LDHB^-/-^ mice at the previously established peak of IFN-I production (3.5 hrs) (Fig. 3d) indicating that LDHB is necessary for optimal pDC IFN-I production *in vivo*.

To determine if LDHB in pDCs contributed to their capacity for IFN-I production during infection, and promoted anti-viral responses, we made mixed BM chimeras using a 50:50 ratio of BDCA2-DTR transgenic and LDHB^-/-^ or WT BM respectively (Fig. 3e). BDCA2-DTR mice express the diphtheria toxin (DT) receptor under the control of the human BDCA2 promoter and allow for the specific deletion of pDCs upon application of DT^60^. Thus, in these chimeras much of the immune compartment is composed of WT cells with normal function^60^, however, after DT treatment only pDCs from the non-DTR fraction (WT or LDHB^-/-^) remain (Extended Data Fig. 3a). Thus, we refer to these animals as pDC-WT and pDC-LDHB^-/-^.

We chose mouse hepatitis virus (MHV) as a model system to investigate the importance of LDHB in pDCs *in vivo*. MHV is a murine specific beta coronavirus and relative of human coronaviruses which cause severe acute respiratory syndrome (SARS-CoV-1, SARS-CoV-2) and middle east respiratory syndrome (MERS). As with SARS-CoV-1^61^, SARS-CoV-2^62^, and MERS^63^, MHV replication is highly dependent on pDC derived IFN-I^64^, and therefore a reasonable model virus to test the *in vivo* relevance of LDHB expression in pDCs.

At day 2 p.i., following depletion with DT we analyzed interferon expression in pDCs, and viral titers in the liver (Fig. 3f,g). In line with our *in vitro* and *in vivo* results with CpG stimulation (Fig. 3b-d) IFN-I production was significantly reduced in pDCs obtained from the pDC-LDHB^-/-^ versus pDC-WT infected mice (Fig. 3f). In conjunction with this we found reduced viral control (greater viral titers) in the livers of pDC-LDHB^-/-^ versus pDC-WT MHV infected mice (Fig. 3g), demonstrating that LDHB in pDCs promoted early MHV control. At this timepoint, we did not observe detectable viral titers in other organs for most mice (Extended Data Fig. 3-c) although we did observe virus in the spleens and lungs of some, but not all pDC-LDHB^-/-^ mice (Extended Data Fig. 3b,c), indicating that while loss of control is not as complete as that observed with total IFNAR deficiency^64^ there may be a tendency toward broader organ infectivity in pDC-LDHB^-/-^ mice. Finally, we did not observe any reduction in CD86 or MHC-II expression in LDHB^-/-^ vs. WT pDCs, and in fact, MHC-II appeared to be increased in pDCs from pDC-LDHB^-/-^ mixed BM chimeras (Extended Data Fig. 3d). This small increase is, however, difficult to disentangle from the higher viral titers observed in pDC-LDHB^-/-^ mice (Fig. 3g), but generally supports that LDHB is not necessary for pDCs expression of antigen presentation molecules after infection. In contrast, our data show that LDHB expression in pDCs is essential for their optimal IFN-I production and early viral control after a beta coronavirus infection.

### LDHB is essential to human pDC function and downregulated in HIV infected patients

Our above experiments describe a role for LDHB in the support of optimal IFN-I production in mouse pDCs. However, there are species-specific differences in pDCs from humans and mice^65^. Thus, we assessed whether LDHB was necessary for human pDC IFN-I production. For this, we made use of Cas9 ribonucleoprotein (RNP) technology. Briefly, RNP were assembled with a fluorescently tagged structural RNA and a crRNA targeting LDHB or a non-targeting control crRNA. These complexes were transfected into purified human pDCs; 24 hrs later these pDCs were stimulated with R848 and analyzed for cytokine production (Fig. 4a-c). Analysis of genome editing by T7 assay revealed low but readily detectable (∼14%-33%) editing (Extended Data Fig. 4a) in line with our results showing modest but significantly reduced LDHB protein expression by flow cytometry (Extended Data Fig. 4b,c). Our results showed that pDCs receiving LDHB targeting gRNAs produced less IFNα than those receiving control gRNA (Fig. 4b,c), but notably had no deficiency in TNFα production (Extended Data Fig. 4d) in line with our results from genetically deficient mice (Extended Data Fig. 2j) suggesting that genetic deficiency in LDHB specifically reduces IFN-I production.

**Figure 4.**
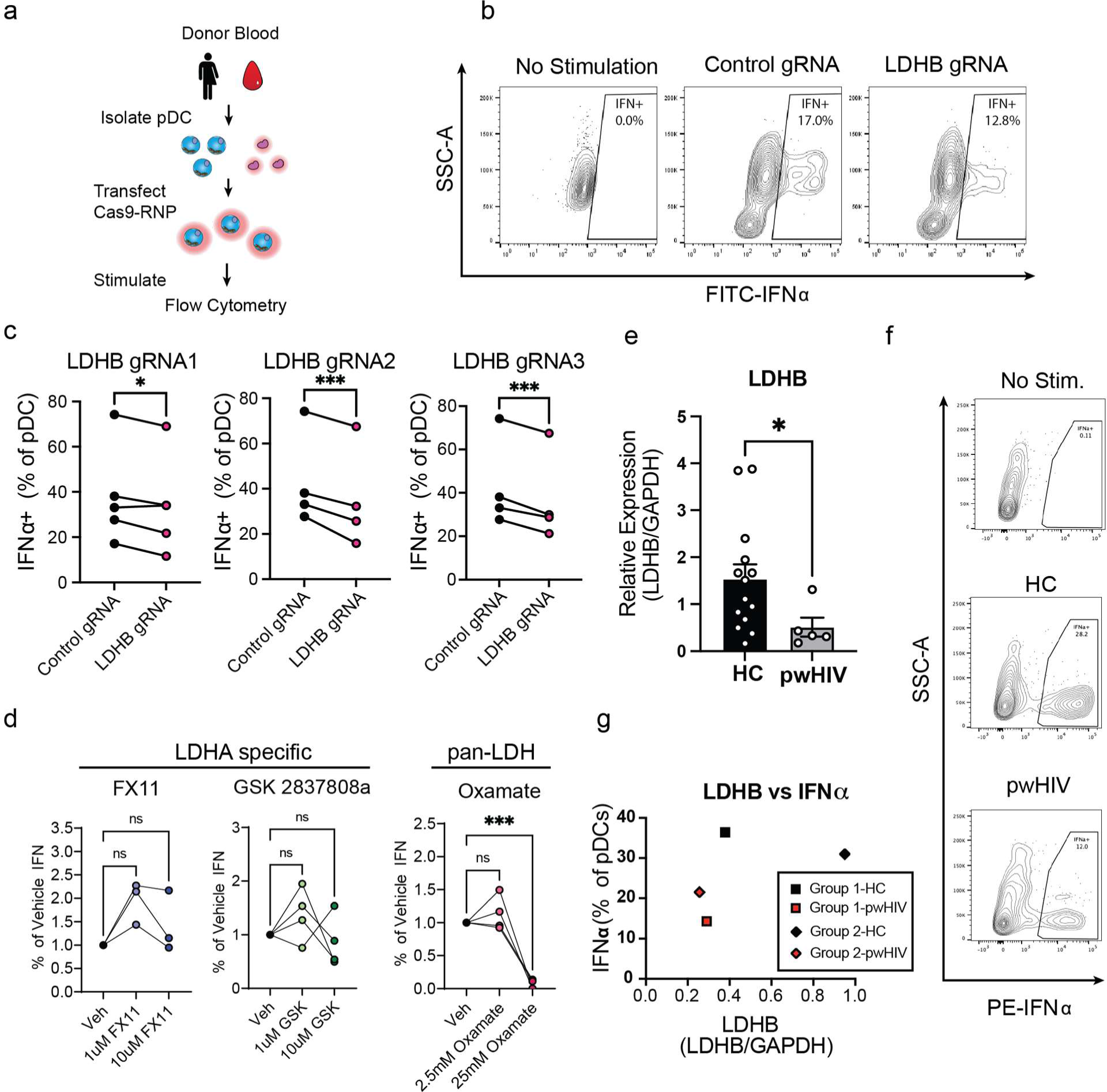
Human pDCs require LDHB for optimal IFN-I production. **(a)** Experimental outline for the modulation of gene expression in primary human pDCs. Breifly, PBMCs were isolated from donor blood, and pDCs were purified, then transfected with Cas9-RNP containing non- targeting control or targeting LDHB. 24 hrs later pDCs were stimulated with R848 and cytokine production measured by flow cytometry. **(b)** Representative plot of IFNα expression in Cas9-RNP transfected human pDCs after stimulation with R848. **(c)** IFNα production in Cas9-RNP transfected human pDCs. **(d)** Human PBMC were isolated from healthy donors, treated with the indicated small molecule inhibitors for 2 hrs prior and concurrent with stimulation, and IFNα was measured in pDCs by flow cytometry 8 hrs after stimulation with CpG-A. (**e**) qPCR quantification of LDHB transcripts in pDCs isolated from healthy controls (HC) or people with HIV (pwHIV). (**f**) Representative plot of IFNα staining from unstimulated HC (up), and HC, pwHIV stimulated with HSV-1 (middle and bottom). (**g**) comparison of LDHB expression and percentage of IFNα producing pDCs in two pairs (group 1 and 2) of HC and pwHIV processed in parallel. Data are representative of 4-5 independent experiments with unique donors **(c)** pooled from 3-5 independent experiments with unique donors **(d),** pooled averages from 19 patients quantified across 3 independent experiments **(e)**, or 2 groups of age/sex matched HC and pwHIV (**g**). Data are shown as mean ± SEM. Statistics used are Paired T Tests **(c),** Unpaired T test with Welch’s correction **(e)** or One-Way ANOVA with Dunnett correction **(d)**. p<0.05 *, p<0.01 **, p<0.001 ***.

To coordinately validate these results with a pharmacological approach we made use of available LDH inhibitors. As an LDHB specific inhibitor with validated capacity to inhibit LDHB activity *in vivo* has not been reported, we made use of Oxamate (OA) an LDH inhibitor which targets both LDHA and LDHB equally^66,67^ and contrasted this with inhibitors specific for LDHA (GSK2837808a^68^, or FX11^69^). We treated at a variety of concentrations within or below previously described EC50 ranges for each drug (GSK2837808a,11 0.4-11uM^68,70^ ; FX11, 49-60uM^71^; OA, 45-76mM^72^) and including the highest tested concentration that did not affect pDC viability (Extended Data Fig. 4e). Treatment with the selective LDHA specific inhibitors GSK2837808a or FX11 had no impact on IFN-I production in pDC from healthy donors after stimulation with CpG-A (Fig. 4d). In contrast, OA treatment severely reduced pDC capacity for IFN-I production (Fig. 4d) without compromising pDC viability (Extended Data Fig. 4e). TNFα production was compromised after OA treatment, suggesting a requirement of LDH for TNFα production in this case (Extended Data Fig. 4f). Similarly, MHC-II expression was reduced with OA treatment (Extended Data. Fig. 4g). Notably, these results match what has been previously reported for GSK2837808a^50^ and suggest that activity of LDHA, but not LDHB, is dispensable for optimal human pDC IFN-I production after CpG-A stimulation.

Like mice, human pDCs show reduced capacity to produce IFN-I in the face of chronic viral infections such as HIV^73–76^ HCV^77–81^, and HBV^82–84^. To determine if LDHB downregulation is a conserved feature of pDC exhaustion we investigated the expression of LDHB in pDCs from people with HIV (pwHIV) who have significantly reduced numbers of pDCs^73–75,85–87^. Purity of isolated pDCs was not different across groups (Extended Data Fig. 4h, Extended Data Fig. 5), with total purity ranging from 90-99% (Extended Data Fig. 4h, Extended Data Fig. 5, Supplementary Table 10). In agreement with our data from mice, LDHB expression was significantly decreased in pDCs from the peripheral blood of pwHIV specifically those with viremia (Fig. 4e). We also investigated whether LDHB expression related to the function of pDCs from pwHIV. pwHIV have significantly reduced numbers of pDCs. Thus, due to extremely limited available material, we were only able to make this comparison for two pairs of donors. Comparison of stimulation of pwHIV and age/sex matched healthy controls revealed that these two patients were deficient in both LDHB expression and IFN production compared to their counterparts (Fig. 4f-g). These data are in line with a model in which LDHB expression and IFN-I production are both suppressed in pwHIV, though of course this must be interpreted with the caveat that this is a low number of samples. Altogether our results using genetic deletion, chemical inhibition, and investigation of LDHB in pwHIV support the assertion that LDHB contributes to optimal production of IFN-I by human pDCs, and LDHB downregulation is a conserved hallmark of exhausted pDCs in both humans and mice.

### pDCs and cDC2 show distinct metabolite preferences, and LDHB is essential to maintain optimal pDC metabolic capacity

Given that LDHB is uniquely expressed in pDCs we hypothesized that there may be differences in energetic carbon sources used by pDCs as compared to conventional DCs. To investigate this, we generated pDCs or cDC2 by Flt3L culture, purified these cells by FACS and incubated them with either ^13^C labeled lactate, or ^13^C labeled glucose then measured incorporation into glycolysis and TCA related metabolites (Fig. 5a). Unexpectedly, we found that pDCs and cDCs similarly incorporated ^13^C from exogenously added lactate into their TCA cycle related metabolites (Fig. 5b). However, we observed a significantly reduced proportion of pDC TCA metabolites derived from glucose (Fig. 5c), suggesting that pDCs make less use of glucose (relative to lactate) to fuel their TCA cycles than cDC2 do comparatively under these conditions (Fig. 5c). Interestingly, concurrent stimulation from the initiation of culture with CpG did not alter lactate or glucose usage by either cDC2 or pDCs (Extended Data Fig. 6a-d).

**Figure 5.**
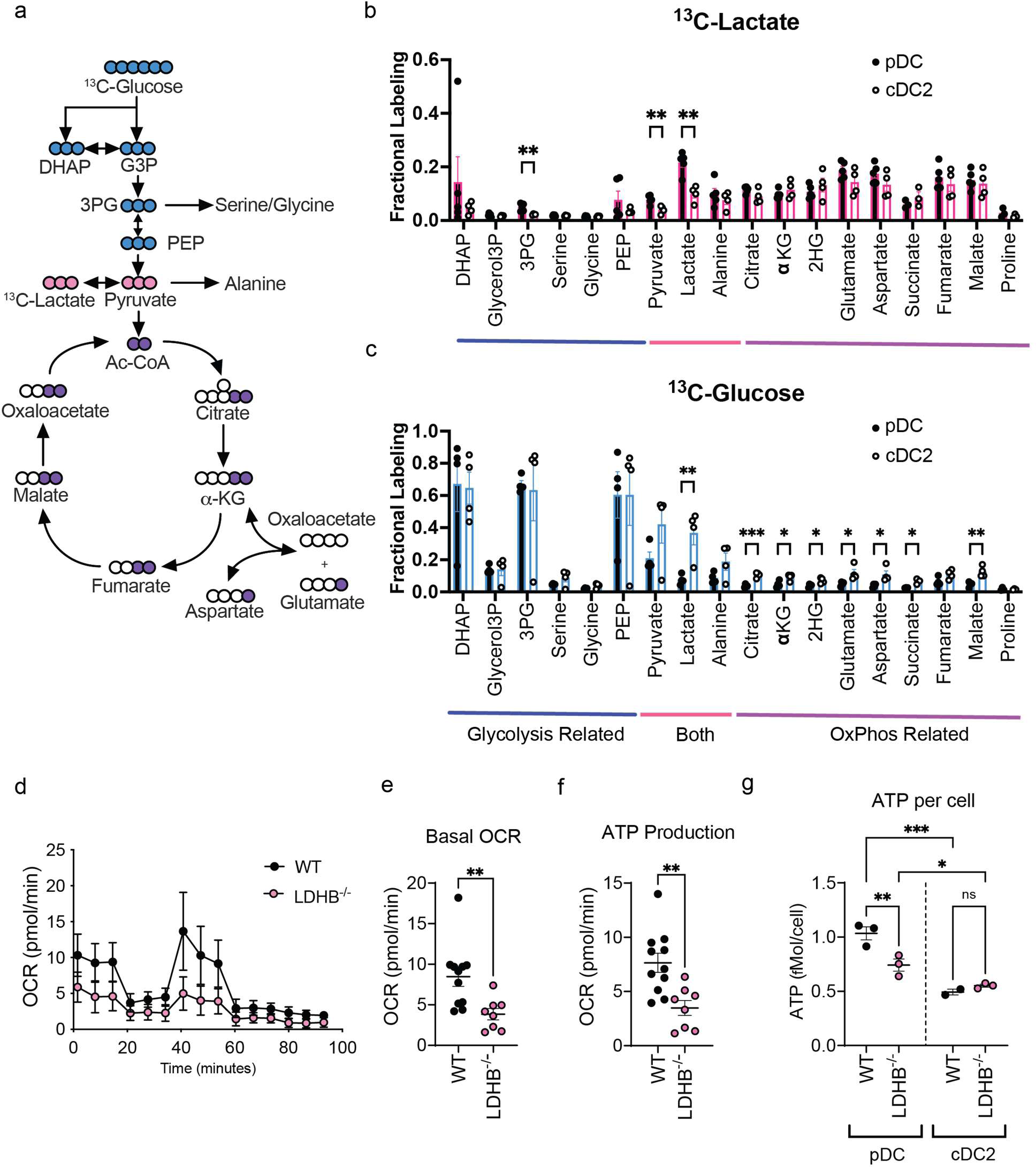
Both glucose and lactate feed the TCA cycle in DCs but pDCs are less labeled from glucose. **(a)** Diagram of stable-isotope tracing incorporation of ^13^C-Lactate and ^13^C-Glucose into glycolytic and TCA intermediates in pDCs isolated from Flt3L culture by FACS at day 8 p.c. **(b,c)** Fractional labeling of glycolysis and TCA cycle intermediates in pDCs and cDC2 by ^13^C-Lactate **(b)** and ^13^C-Glucose **(c)** glycolysis and TCA cycle related metabolites are indicated below the graphs. **(d) S**eahorse tracing of WT (black) or LDHB^-/-^ (pink) pDCs. **(e,f)** Derivative measures from Seahorse assay of WT (black) or LDHB^-/-^ (pink) pDCs **(e)** Basal OCR **(f)** ATP production. **(g)** Total ATP levels in WT (black) or LDHB^-/-^ (pink) pDCs or cDC2. Data are pooled from 2-4 independent experiments **(a-f)** or representative of 2 independent experiments (**g)**. Data are shown as mean ± SEM. Statistics used are Student’s T Tests **(b,c,e,f)** or Two-Way ANOVA with Fisher’s LSD **(g)**. p<0.05 *, p<0.01 **, p<0.001 ***.

We then attempted to determine if LDHB deficiency in pDCs modified their metabolic capacity. Indeed, we found that LDHB deficient pDCs showed significantly reduced basal OCR, and ATP production (Fig. 5d-g), but not reduced basal glycolysis (as defined by ECAR) (Extended Data Fig. 6e,f), or SRC (Extended Data. Fig. 5g). Concurrently, we observed that LDHB deficient pDCs but not cDC2 showed reduced levels of ATP (Fig. 5g), in line with our observations that exhausted pDCs exhibit low ATP content (Fig. 2i). Overall, these data characterize the differential carbon source usage of lactate and glucose in pDCs versus cDC2s, and demonstrate that pDCs and cDC2s can both use significant amounts of lactate and glucose as carbon sources to fuel their oxidative metabolism, and LDHB deficient pDCs show significant defects in oxidative metabolism and ATP content.

### Restoration of LDHB rescues functionally exhausted pDCs

Given the essential role for LDHB in optimal pDC IFN-I production combined with its downregulation in exhausted pDCs, we then sought to determine if restoration of LDHB to exhausted pDCs could rescue their function. For this we harvested BM cells from mice at day 30 p.i. and transduced them with a vector expressing LDHB in combination with enhanced (e)GFP or vector control at day 2 & 3 post-culture (p.c.) with Flt3L. At day 8 p.c., we stimulated cells with CpG-A and analyzed cytokine expression in transduced pDCs. We observed a significant increase in IFN-I production in LDHB transduced cells relative to those transduced with vector control, though this was at the lower end of the range of levels of IFN-I produced by pDC derived from the BM of uninfected animals (Fig. 6a). Along with our data indicating that LDHB deficiency does not completely ablate pDC IFN-I production or change TNFα production capacity these data reinforce the idea that multiple mechanisms are likely at play in the enforcement of pDC exhaustion. Still, it is important to note that this is the first identified method that can restore any amount of exhausted pDC function.

**Figure 6.**
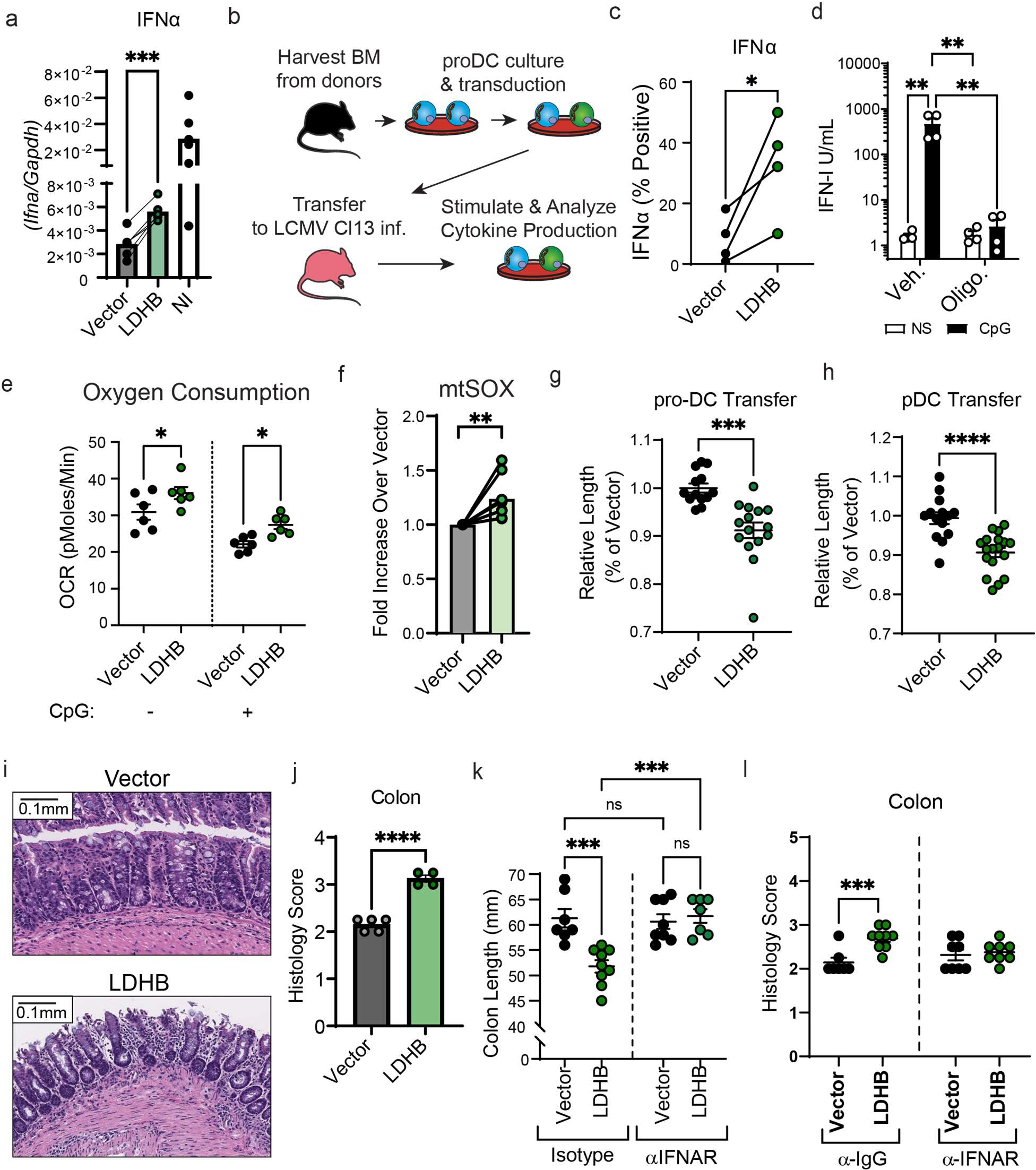
Enforced LDHB expression in exhausted pDCs restores function, and associates with potentiated infection-induced pathology. **(a)** Bone marrow from LCMV Cl13 infected mice at day 30 p.i. was isolated and subjected to Flt3L culture. Cultures from individual mice were separated and half-each transduced with retrovirus encoding LDHB or vector control. At day 8 p.c., pDCs were isolated by FACS and stimulated for 12 hrs with CpG and *Ifna* transcript was analyzed by qPCR. **(b)** Outline of proDC transfer experimental design to analyze pDC functional restoration *in vivo*. Bone marrow from uninfected mice was subjected to Flt3L culture and transduced with retrovirus encoding LDHB or vector control. At day 3.5 p.c. pro-DC were isolated by FACS and transferred into LCMV Cl13 infected mice at day 7.5 p.i. 6 days later splenocytes were isolated from recipients, and subjected to stimulation for 8 hrs with CpG-A, then interferon production in pDCs was assessed by flow cytometry. **(c)** IFNα production in pDCs derived from pro-DCs with enforced expression of vector control or LDHB was measured by flow cytometry. **(d)** pDCs isolated from the spleens of non-infected mice were treated with 1µM Oligomycin for 2 hrs then stimulated with CpG for 6 hrs and IFN-I was measured by bioassay. **(e,f)** Flt3L-culture- derived pDCs from day-30 LCMV Cl13 infected mice, expressing vector control or LDHB, were analyzed for OCR via Seahorse Xfe96 prior to or 30 minutes after stimulation with CpG **(e)** or mitochondrial derived superoxide **(f)**. **(g)** Colon length was measured in LCMV Cl13 infected mice transfered with pro-DCs with enforced expression of vector control or LDHB, as depicted in b, at day 14 p.i. (day 6 post transfer). **(h)** Colon length was measured in LCMV Cl13 infected mice transferred with differentiated Flt3L-derived pDCs, with enforced expression of vector control or LDHB, at day 11.5 p.i. (day 2.5 post transfer). **(i,j)** Representative **(i)** and quantitative **(j)** analysis of colon histology from LCMV Cl13 infected mice receiving differentiated pDCs as described in g. **(k,l)** Colon length **(k)** and histology score **(l)** were depicted as described in g for mice receiving differentiated pDCs and concurrently treated with either Isotype control antibody or IFNAR blocking antibody. Measurements were made blinded (**g,h,j,k,l)**. Data are representative of 2 **(a,i,j)** experiments **(a)**, or pooled from 2 **(d,e,k,l)**, 4 **(c,f,g)**, or 5 **(h)** independent experiments. Data are shown as mean ± SEM. Statistics used were paired T Test **(a,c,f)** One-way ANOVA with Tukey Correction **(d)**, Student’s T Test **(e,g,j),** or two-way ANOVA with Fisher’s LSD **(k,l)** p<0.05 *, p<0.01; **, p<0.001 ***, p<0.0001 ****.

To further investigate whether LDHB restoration recovers pDC cytokine production *in vivo,* we made use of a newly developed system of pro-DC transfer. Briefly, *in vitro* developed pro-DCs, which can generate cDCs & pDCs^88^, were transduced to modulate gene expression, then transferred *in vivo* to complete their development into mature DCs (Fig. 6b). BM cells from uninfected mice were cultured with Flt3L and transduced at days 2-3 p.c. pro-DCs were then isolated at day 3.5 p.c. and transferred into LCMV Cl13 infected mice at day 7.5 p.i. Six days later (day 13.5 p.i.) we isolated splenocytes from these mice, stimulated them with CpG-A, and assessed the IFNα production of pDCs developed from the transferred pro-DCs by flow cytometry. We found consistently that LDHB overexpressing pDCs indeed had greater production of IFNα after isolation from infected mice (Fig. 6c). Together these data demonstrate that LDHB restoration can promote IFN-I production capacity in exhausted pDCs *in vitro* and *in vivo*.

### pDCs require oxidative metabolism for optimal IFN-I production and LDHB restoration improves oxidative metabolism in exhausted pDCs

It has recently been shown that oxidative metabolism is essential to human pDC function^50^ to directly assess whether this is also true in mice we isolated pDCs from naïve animals, and pretreated them with Oligomycin (an inhibitor of ATP synthase), then stimulated them with CpG and measured IFN-I production by bioassay (Fig. 6d). This treatment completely ablated pDC IFN-I production, demonstrating that like human pDCs, mouse pDCs require oxidative metabolism for their production of IFN-I. We then investigated how LDHB restoration might improve the sustained metabolic deficiencies observed in exhausted pDCs. For that, we first measured oxygen consumption in exhausted pDCs expressing LDHB or vector control, generated as in Fig. 6a. We found that pDCs expressing LDHB showed greater levels of oxygen consumption, and better sustained oxygen consumption following stimulation with CpG (Fig. 6e). In line with these data, expression of LDHB in exhausted pDCs associated with significantly greater quantities of mtSOX, indicating that LDHB enforcement may improve oxidative metabolism in exhausted pDCs (Fig. 6f). Altogether these results suggest that pDCs require oxidative metabolism for IFN-I production, and enforcing LDHB expression in exhausted pDCs can improve oxygen consumption.

### LDHB expression in pDCs enhances virus-associated pathology

Despite exposing hosts to increased risk of secondary infection, pDC exhaustion appears to be conserved across LCMV infection in mice^3,4,18^, SIV infection in macaques^89,90^, and humans infected with HIV^73–76^, HCV^77–80,91^, HBV^82–84^ or SARS-CoV-2^92^. This conservation suggests that pDC exhaustion is an important feature of pDC biology, but its purpose is unknown. One salient but untested hypothesis is that pDC exhaustion, like CD8 T cell exhaustion^93^, exists to protect the host from overt immune driven pathology. To investigate this, we returned to the proDC transfer model that allowed pDC functional restoration (Fig. 6b,c). LCMV Cl13 infection causes colitis-like symptoms including colon shortening^94^, additionally in non-infectious conditions, release of pDC negative regulators has been established to drive intestinal pathology^95^. Therefore, we focused our investigation on the intestinal compartment of these mice. Intriguingly, in 4 of 4 experiments we observed shortening of the colons for mice receiving LDHB expressing proDCs as compared to their vector transferred counterparts (Fig. 6g). Notably, while IFN-I is critical for many viral infections^1^ treatment with recombinant IFNα is not impactful for viral replication at later stages after LCMV Cl13 infection^96^. Accordingly, we observed only a mild reduction in viral titers that did not reach significance when transferring LDHB expressing proDCs in LCMV Cl13 infected mice (Extended Data Fig. 7a).

It is important to recognize that proDCs can differentiate into both cDC1 and cDC2 as well as pDC. Although we did not observe any alterations to cDC proportion, cDC function, or DC maturation (as measured by expression of CD86) in the populations derived from LDHB expressing proDCs after transfer (Extended Data Fig. 7b-d), suggesting that cDCs are similar in both conditions, these experiments did not directly demonstrate a relationship between LDHB restoration in pDC and colitis. Thus, to determine whether the enhanced colon shortening was pDC dependent, we performed similar experiments transferring mature Flt3L derived pDCs expressing LDHB or vector control at day 8.5 p.i. and investigating colon length 2.5 days later at day 11 p.i.. As expected, mice receiving LDHB expressing pDCs exhibited shorter colons than their vector control counterparts (Fig. 6h). Furthermore, histological examination found increased inflammation in these same colons (Fig. 6i,j) confirming that transfer of LDHB expressing pDCs into infected mice associated with increased colitis. Intriguingly, we observed no significant increase in pathology in the small intestine or liver of these animals (Extended Data Fig. 7e,f), suggesting that some tissues may be more susceptible to pathology caused by LDHB-expressing pDCs during viral infection.

We then sought to determine whether this increased inflammation was the result of IFN-I production. To this end, we repeated the experiments as described, but provided mice transferred with vector or LDHB with either IFNAR blocking antibody^97^ or isotype control antibody immediately following the transfer of vector or LDHB expressing pDCs, and again at days 0.5 and 1.5 post-transfer. The mice treated with isotype control antibody recapitulated what we observed in untreated mice, showing shorter colons in the group transferred with LDHB expressing pDCs versus their vehicle control group (Fig. 6k). In contrast, treatment with anti-IFNAR resulted in full restoration of colon length in the mice with LDHB expressing pDCs (Fig. 6k), and eliminated the difference in both colon length and inflammatory score by histology between animals transferred with LDHB-expressing or vector control pDCs (Fig. 6k&l). Altogether these results demonstrate that LDHB expression in pDCs restores exhausted pDC function, and that during *in vivo* viral infection, the presence of pDCs that resist exhaustion was accompanied with increased IFN-I dependent colitis. This suggests that LDHB expression and pDC exhaustion manage a trade-off between supporting pDC function and opposing host-pathology during viral infection.

## Discussion

Type I IFN is a critical antiviral and antineoplastic cytokine. As pDCs produce the largest quantities and most types of IFN-I of any cell type, they are critical first responders for many viral infections^2^. Accordingly, pDC exhaustion associates with increased susceptibility to secondary infection^3,4,73^. Until now there has been no method to restore exhausted pDC IFN production, and so there has been no tool available to understand the physiological consequences of ablating this well conserved process^3,4,73–79,81–84,89,90^. Our work has established that pDC exhaustion is associated with profound metabolic deficiencies, and that LDHB supports the metabolism of pDCs as well as the production of IFN-I by pDCs in both humans and mice. We showed that LDHB expression in pDCs was suppressed by persistent infections and that restoration of LDHB reinstated IFN-I production capacity in the otherwise exhausted pDCs *in vitro* and *in vivo*. Furthermore, this allowed us, for the first time, to determine the consequence of restoring function to exhausted pDCs. In the context of persistent LCMV infection, enforced LDHB expression enhanced virus-induced-colitis^94^ in an IFNAR dependent manner, suggesting that pDC exhaustion may be a conserved mechanism which provides protection against excessive pathology during viral infection.

Through our analysis we generated the first gene signature of pDC exhaustion. Notably, this independently identified several known regulators of pDC IFN-I production (*Cd81*^45,46^*, Myc*^47^) as well as several genes with unknown relationship to pDC function, but previously identified as differentially expressed in pDC from SIV-infected macaques (*Tigit*)^44^. Tigit is of particular interest since it is a surface expressed protein and a known regulator of exhaustion in CD8 T cells and being explored as a target in cancer immunotherapy^98^.

Our study identified *Ldhb* as a “pDC exhaustion signature” gene that is downregulated after infection while in uninfected mice was uniquely highly expressed in pDCs as compared to other DCs. This may be because LDHB is a target of E2-2 (encoded by *Tcf4*), an essential transcriptional regulator of the pDC lineage^99,100^ which we have previously shown to be downregulated in exhausted pDCs from both humans and mice^18^. It is thus tempting to speculate that reduced levels of E2-2 drive LDHB downregulation in exhausted pDCs, however further work will be needed to support this hypothesis.

By identifying LDHB as an essential regulator of pDC IFN-I production, our study adds to a significant body of work by other groups understanding essential metabolic regulators of pDC function at steady state, or shortly following stimulation ^48–50,101,102^. Our study does much to reinforce general principals discovered by these groups, including an essential role for oxidative metabolism in pDC IFN-I production ^48–50,101^. Importantly, our study is the first to investigate how pDC metabolism changes *in vivo* following infection, and how these metabolic adaptations relate to the development of pDC exhaustion. It is important to address that it was previously reported that LDH activity is dispensable for human pDC IFN-I production^50^, the inhibitor used in this case (GSK2837808a) has high specificity for LDHA, and so this is still in line with our observation that human and mouse pDCs rely on LDHB, but not LDHA, for IFN-I production. Indeed, we were able to recapitulate the fact that GSK2837808a does not impact pDC IFN-I production after CpG- A simulation, reinforcing the dispensability of LDHA specifically in regulating pDC IFN-I production (Fig. 4d).

Our work also demonstrates for the first time that both pDCs and cDCs can use extracellular lactate as a significant carbon source for TCA cycle metabolites on par with glucose. This is a potentially important observation as, in the context of inflammatory environments, where other cells such as activated macrophages, T cells, or tumor cells (in the case of a tumor microenvironment) consume large quantities of glucose and excrete large quantities of lactate, DCs may need to rely on lactate to support oxidative metabolism. Further work will be needed to pursue this hypothesis, as well as to determine the preferred carbon sources of these cells in supporting their TCA cycles including assessment of glutamate and fatty acid oxidation.

Very little is understood about pDC IFN-I exhaustion in the context of cancer, but it has been described across a variety of types of malignancies^21–24^, suggesting the phenomenon is at least somewhat conserved. Adjuvant therapies that stimulate pDCs to produce IFN-I have shown great promise in the treatment of some varieties of cancer^25–28^, and our study suggests that coordinated metabolic and stimulatory intervention may be more successful than either approach alone. On the other hand, our work also provides insight into what types of therapies may inadvertently compromise pDC activity. A generalized LDH inhibition, for example, has been proposed as a mechanism to treat highly glycolytic cancers. Our study suggests that inhibitors which do not target LDHB may be more effective as they are likely to avoid further suppressing pDC function.

Given the well-established anti-viral and antitumor roles of IFN-I^1,13^, it has previously been unclear why pDC exhaustion is conserved across species. Our study provides one possible reason in that we observe more severe virus associated pathology in animals where pDC exhaustion is opposed. In addition, following an infection, it may also be beneficial to default to the active suppression of IFN-I production as there are several bacterial infections, including ancient infections (e.g., *Mycobacterium tuberculosis, Listeria monocytogenes*, *Staphylococcus aureus*), for which IFN-I responses can be detrimental^103–106^. Furthermore, as most viruses encode highly effective interferon evasion strategies ^107^ continued production of IFN-I after establishment of infection may have diminishing returns for viral control. Indeed, treatment with recombinant IFN- β and IFN-α5 at late stages of LCMV Cl13 infection did not affect viral titers^96^ and persistent interferon signaling counterintuitively leads to CD8 T cell exhaustion and promotes viral persistence^108,109^. Conversely, early application of IFN-β and IFN-α5 in the same infection model leads to improved viral control^96^, therefore delayed pDC exhaustion might be expected to improve viral control, although the relative tradeoff in immunopathology in this case is not known. Interestingly, pDC functional impairment has been described in systemic lupus erythematosus (SLE), a fact sometimes used to assert their lack of importance for this disease^110^. However, it is possible that pDC exhaustion emerges after transient IFN-I production (as in the viral infection), that de novo generated pDCs may continuously provide a new source of IFN-I prior to becoming exhausted. Additionally, it is possible that pDC exhaustion is incomplete in these patients as no directly comparable assessment of pDC exhaustion capacity, such as a comparison of virally induced pDC exhaustion in these patients prior to developing SLE, has been evaluated.

Altogether, our study characterizes the transcriptional state of pDCs throughout infection, and as a result identifies large scale metabolic reprogramming in pDCs following viral infection. We identify LDHB as a novel regulator of human and mouse pDC IFN-I production which is downregulated during pDC exhaustion and demonstrates that restoring LDHB rescues IFN-I production capacity to exhausted pDCs. This is associated with enhanced IFNAR dependent infection-induced-colitis when LDHB expressing pDCs are transferred into infected mice, suggesting the conservation of pDC exhaustion may be a mechanism to avoid excessive pathology during infection. These data provide the first proof-of-concept showing that it is possible to restore exhausted pDC function, and the first evidence that the purpose of conservation for pDC exhaustion across species is to protect against immunopathology. Furthermore, our study provides insights into the metabolic machinations pDCs use to maintain optimal IFN-I production, and concurrently identify novel avenues to manipulate pDC function for the benefit of human health.

## Methods

### Mice

Six to eight-week-old female C57BL/6 and B6 CD45.1 mice were purchased from The Jackson Laboratory. TLR7^-/-^ mice and BDCA2-DTR mice were purchased from The Jackson laboratory and bred in house. LDHB^-/-^ mice were generated by the international mouse phenotyping consortium (IMPC, www.mousephenotype.org) and procured from the Knockout Mouse Consortium (KOMP). All mice were housed under specific-pathogen-free conditions at the University of California, San Diego. For pDC depletion mixed BM chimeras of 50% BDCA2- DTR and 50% either WT or LDHB^-/-^ BM were treated with diphtheria toxin (DT) (Sigma-Aldrich, D0564) as described^60^. Briefly, mice were given 200 ng of DT by intraperitoneal injection daily starting from two days prior to MHV infection, and up to endpoint (48 hrs p.i.). Mouse handling and experiments conformed to the requirements of the National Institute of Health and the Institutional Animal Care and Use Guidelines of UC San Diego. Unless stated otherwise, experiments were initiated in mice (female and male) at 7-12 weeks of age.

### Virus strains

LCMV Infection was performed by injecting mice with 2 × 10^6^ plaque-forming units (pfu) of LCMV Cl13 intravenously (i.v.) via the tail vein. LCMV Cl13 was propagated in BHK cells and quantified by plaque assay performed on Vero cells^111^. Briefly, Vero cell monolayers were infected with serially diluted viral stock and incubated for 60 minutes at 37°C in 5% CO2 with gentle shaking every 15 minutes. Agarose overlay was added to infected cells and placed in an incubator for 6 days at 37°C in 5% CO2. Cells were fixed with formaldehyde and stained with crystal violet for 5 minutes at room temperature and plaques were counted. MHV A59 was a generous gift from Prof. Susan Weiss. MHV infection was performed by injecting 50 p.f.u./mouse intraperitoneal (i.p.). MHV titers were assessed as described above for LCMV and as reported previously^112^, using L929 cells from ATCC (ATCC: CCL-1) and incubating with agar overlays for 2 days prior to formaldehyde fixation.

### Human Samples Inclusion and Ethics

All human subject studies were approved by the Institutional Review Board (IRB) at Rutgers, the State University of New Jersey, New Jersey Medical School. Blood samples from HIV-negative and HIV-infected volunteers were obtained with consent according to institutional guidelines and the Declaration of Helsinki.

### Human pDC Isolation

Human pDC isolation was performed using an EasySep™ Human Plasmacytoid DC Enrichment kit (STEMCELL Technologies) following the manufacturer’s instructions.

### Cas9 RNP Generation and Transfection into human pDCs

Alt-R® Cas9-guides were predesigned by integrated DNA technologies (IDT), sequences reported in Supplementary Table 11. Cas9 RNP were generated using Alt-R® S.p. HiFi Cas9 Nuclease (IDT), Alt-R® CRISPR-Cas9 tracrRNA, ATTO™ (IDT), in combination with control crRNA (IDT) or crRNA targeting LDHB (IDT). Human pDCs were isolated as above and cultured 16 hrs in RPMI with the addition of 10% FCS, 2mM L-Glutamine and 20ng/mL of recombinant human IL-3 (R&D systems). Cas9 and gRNA components were assembled into RNP per the manufacturer’s instructions and transfected into pDC 16 hrs p.c. using Lipofectamine™ CRISPRMAX™ transfection reagent per the manufacturer’s instructions. Cells were then cultured for an additional 24 hrs in the above-described conditions prior to analysis of function.

### Flt3L DC Culture

Flt3L DC culture from infected and uninfected mice was performed as previously described^18,113^. BM DCs were generated at the concentration of 2 x 10^6^ cells/ml for 8 days in 5 ml of DC medium (RPMI-1640 supplemented with 10% (vol/vol) fetal bovine serum, L-glutamine, penicillin-streptomycin, and HEPES pH 7.2) supplemented with 100 ng/mL Flt3L (provided by CellDex Therapeutics) and 50 µM β-mercaptoethanol. Half of the medium was replaced after 5 days with medium with fresh Flt3L added.

### Generation of Retroviral Particles (RV) and Retroviral Transduction

HEK293T cells were transfected with vector or control LDHB, LT1 transfection reagent (Mirus Bio, Madison, WI), and the pCL-ECO packaging plasmid. Supernatants were harvested 48 hrs later and kept at 4°C until media were replaced with fresh media and the second supernatants were harvested 24 hrs after media change (72 hrs post-transfection). Fresh supernatants containing RV without any freezing were used for retroviral transduction. At day 2 post-Flt3L culture, BM cells were transduced with RV encoding vector control or LDHB and 10ug/ml polybrene reagent (Fisher Scientific) and spin-infected at room temperature at 1000g for 90 min. Cells were incubated 37°C overnight. On day 3 post-Flt3L culture, cells were again transduced with RV encoding vector control or LDHB. Cells were then washed in PBS twice and placed in fresh DC medium + Flt3L.

### Cell Sorting

Spleens were incubated with 1 mg/mL collagenase D for 20 minutes at 37°C and pushed through a 100μm strainer to make a single cell suspension. Mouse bones (femur and tibia) were isolated, cleaned of any muscle tissue, then flushed with RPMI with 10% FCS and 2% glutamine to retrieve BM. Splenocytes or BM cells were then subjected to red blood cell lysis in 1 mL ammonium-chloride-potassium (ACK) lysis buffer for 5 minutes. Splenocytes or BM cells were enriched using EasySep™ Mouse Streptavidin RapidSpheres™ Isolation Kit (Stemcell Technologies) per the manufacturer’s instructions. Biotin conjugated anti-Thy1.2 (53-2.1, eBioscience) and anti-CD19 (eBio1D3, eBioscience). Cells were stained with propidium iodide (PI) (Sigma Aldrich) as well as streptavidin conjugated to PerCP-Cy5.5 (eBioscience) (to account for any biotin-conjugated antibody remaining post enrichment) and pDCs FACS-purified by the following definitions: pDC; PI^-^, CD19^-^, Thy1.2^-^, NK1.1^-^, CD11c^+^, CD11b^-^,B220^+^,BST2^+^, cDC1; PI^-^ CD19^-^, Thy1.2^-^, NK1.1^-^, CD11c^+^, CD11b^-^,B220^-^,CD8a^+^, cDC1; PI^-^, CD19^-^, Thy1.2^-^, NK1.1^-^, CD11c^+^, CD11b^+^,B220^-^. Where Flt3L culture was used pDCs were harvested from culture and stained with PI by the following definition: PI^-^, CD11c^+^, CD11b^-^, B220^+^,BST2^+^). proDC^88^ were FACS purified at day 3.5 after Flt3L culture by the following definition: PI^-^, CD19^-^, B220^-^, Thy1.2^-^, CD3^-^, CD4^-^, CD8^-^, CD127^-^, NK1.1^-^, Ter119^-^, Gr-1^-^, MHC-II^-^, CD11c^-^, CD11b^-/low^. All sorting was performed on a BD ARIA II flow cytometer (BD Biosciences).

### Cell Transfer

For proDC and mature pDC transfer cells were FACS isolated at days 3.5 and 8 p.c. respectively following retroviral transduction as described above. For pro-DCs between 2 ×10^5^-5×10^5^ cells were injected intravenously into recipients at day 7.5 p.i. in a total of 200uL in PBS. For mature pDCs between 1 x10^5^-2 x10^5^ cells were injected intravenously into recipients at day 8.5 p.i. and isolated 2.5 days later at day 11 p.i. For anti-IFNAR treatment mice were given anti-IFNAR antibody MAR-1 (500ug/mouse) on the day of transfer, and 250ug/mouse at days 0.5, and 1.5 post transfer by intraperitoneal injection.

### Metabolic Flux Assays

To analyze OCR and ECAR Seahorse Xfe96 plates and cassettes (Agilent) were prepared per the manufacturer’s instructions. On the day of the experiment 1.5×10^5^ pDCs from the BM (primary pDC) or 500,000 pDCs (Flt3L Culture) were isolated by FACS into RPMI Seahorse Assay Medium with the addition of 10% FCS, 2mM L-Glutamine & 10mM Glucose, prior to plating onto Poly-L-Lysine coated 96 well assay plates. Cells were then rested for 1hr at 37C without additional CO2. Assays were either performed with a Seahorse Xfe96 analyzer (Fig. 6e) or a XF HS Mini (All other figures). Mitochondrial stress test assay for pDC on the XF-HS Mini was optimized in-house within the ranges provided by the manufacturer and performed using the mitochondrial stress test kit (Agilent), with the addition of 2-DG (Millipore Sigma, D8375) in the 4^th^ injector port. In well concentrations for each drug at time of injection were Oligomycin (1.5uM), FCCP (2uM), Antimycin/Rotenone (0.5uM) and 2-DG (50mM). Basal OCR was calculated as the difference between OCR prior to any drug addition and OCR after Antimycin/Rotenone. Spare respiratory capacity was calculated as the difference between OCR prior to the addition of any drug and the OCR measured after FCCP addition. ATP production was calculated as the difference between OCR prior to the addition of any drug and the OCR measured after Oligomycin addition. Basal glycolysis was calculated as the difference between ECAR prior to any drug addition and ECAR after the addition of 2-DG.

### ELISA & IFN Bioassay

After FACS purification pDCs were stimulated with 1µM CpG-A (Invivogen ODN 1585) and supernatant was collected for Bioassay or ELISA at 6 or 12 hrs respectively. ELISA and Bioassay were then performed as previously described^18^. For each assay limit of detection was determined as the lowest concentration used for the standard curve multiplied by the dilution factor of the samples, all unstimulated samples were below limit of detection. For stimulated samples below limit of detection values are reported at the limit of detection.

### qRT-PCR

Isolated immune subsets were subjected to RNA extraction using an Rneasy micro kit (Qiagen) including treatment with Dnase I to remove genomic DNA. RNA was reverse transcribed into cDNA using Superscript III RT (Invitrogen). The expression of various genes was quantified using Fast SYBR Green Master Mix (Thermo Fisher Scientific) or TaqMan™ Fast Universal PCR Master mix with probe sets from the Universal Probe Library (Roche). The CFX96 Touch Real-Time PCR Detection System (Bio-Rad) was used to quantify reactions. Primers for the indicated genes are listed in Supplementary Table 12. Relative transcript levels were normalized against mouse *Gapdh* or human *GAPDH* as described previously^18^. Detectable range for each primer set used was determined by evaluating the linear range of a series of dilutions. CT values above detectable range were considered to be non-detected, and their values set as maximum cycle number (45 cycles).

### Flow Cytometry

The following antibodies were used to stain single cell suspensions prepared from murine BM or spleens or mouse Flt3L culture: Thy 1.2 (53-2.1, eBioscience), CD19 (eBio1D3, eBioscience), NK 1.1 (PK136, eBioscience), Gr-1 (RB6-8C5, BioLegend), CD11c (N418, eBioscience), CD11b (M1/70, BD Biosciences), B220 (RA36B2, BioLegend), BST2 (eBio927, eBioscience), CD8 (53-6.7, eBioscience), CD45.1 (A20, eBioscience), CD45.2 (104, eBioscience), MHC class II (I-A/I-E) (M5/114.15.2, BioLegend), CD86 (GL1, eBioscience), Ter-119 (TER-119, eBioscience), CD127 (A7R34, eBioscience), CD3 (145-2C11, eBioscience), CD4 (RM4-5, eBioscience), CD8 (53-6.7, eBioscience). The following antibodies were used to stain human pDCs either from PBMCs or after purification and culture as described in the text: CD3 (UCHT1, BioLegend), CD14 (HCD14, BioLegend), CD16 (3G8, BioLegend), anti-human CD19 (HIB19, BioLegend), anti-human CD56 (MEM-188, BioLegend), HLA-DR (L243, BioLegend), CD11c (B-ly6, BD Biosciences), CD123 (6H6, BioLegend), CD304 (12C2, BioLegend), LDHB (EP1566Y, abcam) TNF-*α* (Mab11, BioLegend), IFN-α (LT27:295, Miltenyi Biotec). PI (Sigma Aldrich) or Ghost dye (Tonbo Biosciences, San Diego, CA) was used to exclude dead cells. Cells were pre-incubated with CD16/CD32 Fc block (BD Pharmingen) for 10 minutes prior to surface staining. Mitochondrial dyes (MitoSOX™ Red Thermo Fisher M36008, MitoTracker™ Red chloromethyl-X-rosamine (CMXRos) Thermo Fisher M46752, MitoTracker™ Green FM M46750) were used per the manufacturer’s instructions. Cells were acquired with a ZE5 flow cytometer (Biorad) and data were analyzed using FlowJo software (BD Biosciences).

### Isotopic Tracing Gas Chromatograph-Mass Spectrometry (GC-MS) Sample Preparation and Analysis

pDCs were isolated by FACS sorting as described above, then rested for 1hr in RPMI with the addition of 10% FCS and 2mM L-Glutamine. Labeled ([3-^13^C]-Lactate, Cambridge Isotope Laboratories CLM-1578) or unlabeled lactate (Sigma-Aldrich) and labeled (U-^13^C6, Cambridge Isotope Laboratories CLM-1396-1) or unlabeled glucose (Thermo Fisher Scientific) were then added to generate a culture with a final concentration of 5mM Lactate and 10mM glucose, and incubated for 3 hrs. Afterward cells were collected, washed in cold 0.9% NaCl, then frozen and stored at -80°C prior to extraction. Metabolites were extracted, analyzed, and quantified as previously described^114^. Mass isotopomer distributions and total metabolite abundances were computed as described previously, labeling is depicted as the fraction of labeled metabolites as described previously^114^.

### Histopathological Analysis

Colon, SI and Liver samples were isolated from mice, fixed, embedded, sectioned, and stained using hematoxylin/eosin as previously described^115^ by the Tissue Technology Shared Resource at UCSD. Scoring of colon and SI inflammation was performed using the inflammation score defined in Erben et al 2014^116^. Scoring of Liver was performed by applied pathology systems (APS) using the method described in reference ^117^.

### Statistics & Reproducibility

Randomization and blinding were used in the measurement of colon length, as well as the scoring of colon inflammation where the measurer was blind to the treatment of the mice. All relevant data are shown as mean ± standard error The number of animals or samples are stated in figure legends. Specific statistical tests are stated in figure legends. We used Student’s t-test when comparing two groups where samples were independently generated, or Paired t-test when samples were derived from the same animal. In cases where variances were unequal, as defined by significant F test, unpaired T test with Welch’s correction was used. Multiple comparisons corrections were used when more than one comparison was performed and are stated in legends. One way ANOVA with Tukey correction was used when there were many independent comparisons. All analyses were performed with Graphpad Prism 10 software.

## Data Availability

RNA-seq and Microarray datasets reported in this study will be uploaded to Gene Expression Omnibus prior to publication. Other data and reagents are available from the corresponding author on request.

## Code Availability

All software used to perform these analyses is publicly available. Software tools are listed in the main text and methods.

## Figure Legends

**Extended Data Figure 1.**
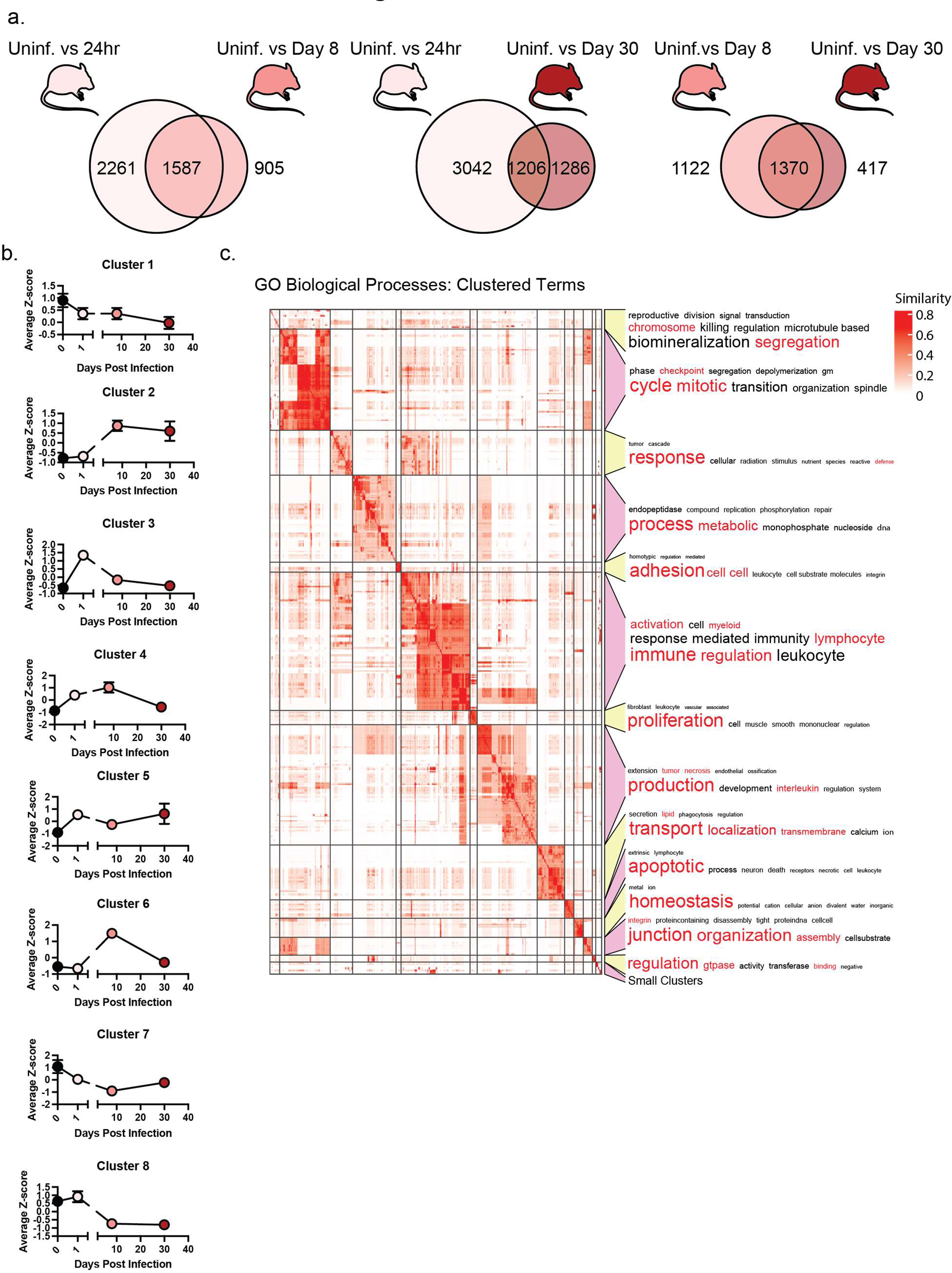
Transcriptional analysis of pDCs throughout persistent viral infection. **(a)** Number of DE or overlapping genes between pDCs isolated from uninfected mice (Uninf.) or from mice at days 1 (24 hr), 8, or 30 p.i. **(b)** Average Z-score for expression of genes in each cluster (as identified in Fig. 1) at each timepoint. **(c)** GO Biological processes identified as enriched in genes differentially expressed both when comparing uninfected to day 8 p.i., as well as comparing uninfected to day 30 p.i. (pDC exhaustion related) were grouped by semantic similarity. Word clouds for each cluster identified by this analysis are presented with larger font representing higher frequency within the biological processes in that cluster.

**Extended Data Figure 2.**
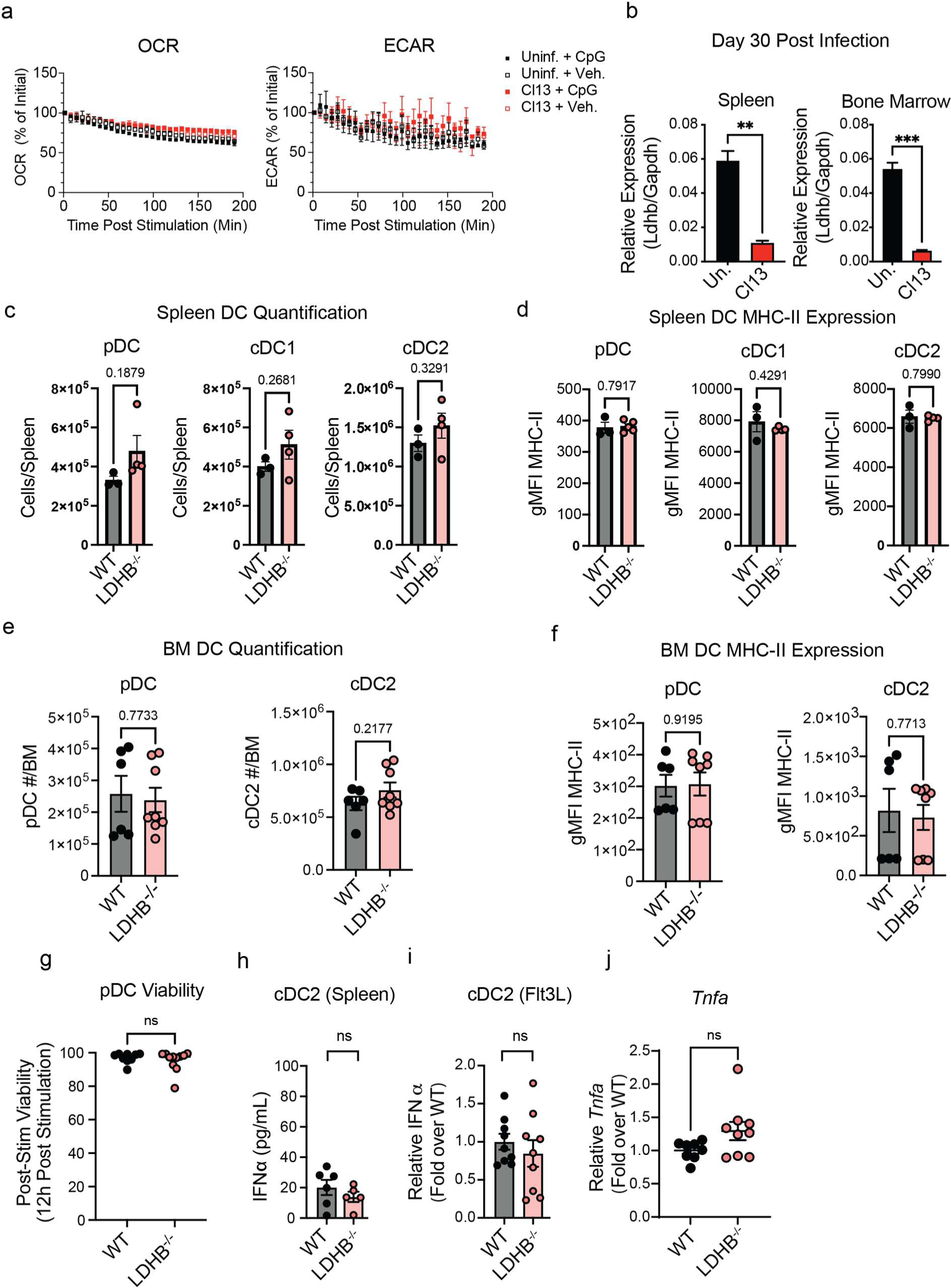
pDCs modify metabolism and expression of LDHB in infection, and LDHB deficiency does not change DC numbers, MHC-II expression, viability or IFN production by cDC2. **(a)** OCR and ECAR traces of pDCs from uninfected mice (black) or LCMV Cl13 infected mice (red) at day 8 p.i. following stimulation with CpG (closed symbols), or no stimulation (open symbols). **(b)** qPCR analysis of *Ldhb* expression in pDCs isolated from the spleen or BM of mice at day 30 p.i. **(c-f).** Number of pDCs **(c,e)**, and expression of MHC-II **(d,f)** were measured by flow cytometry in the spleen (**c,d**) and BM (**e,f**) from WT (black) or LDHB^-/-^ (pink) mice. **(g)** FACS purified pDCs were stimulated with CpG-A for 12 hrs, supernatant collected for ELISA as in Fig. 3b, and viability was measured 12 hrs after CpG stimulation by flow cytometry. **(h,i)** ELISA of IFNα from cDC2 isolated from spleens **(h)** or Flt3L cultures **(i)** from WT (black) or LDHB^-/-^ (pink) mice. **(j)** qPCR analysis of *Tnfa* transcript in Flt3L culture derived pDC stimulated with CpG-A for 12 hrs. Data are pooled from 2-3 **(a,e,f,g-j)** or representative of 2-4 independent experiments **(b-d)**. Data are shown as mean ± SEM. Statistics used Student’s T Test **(b-j)** p<0.05 *, p<0.01; **, p<0.001 ***, p<0.0001 ****.

**Extended Data Figure 3.**
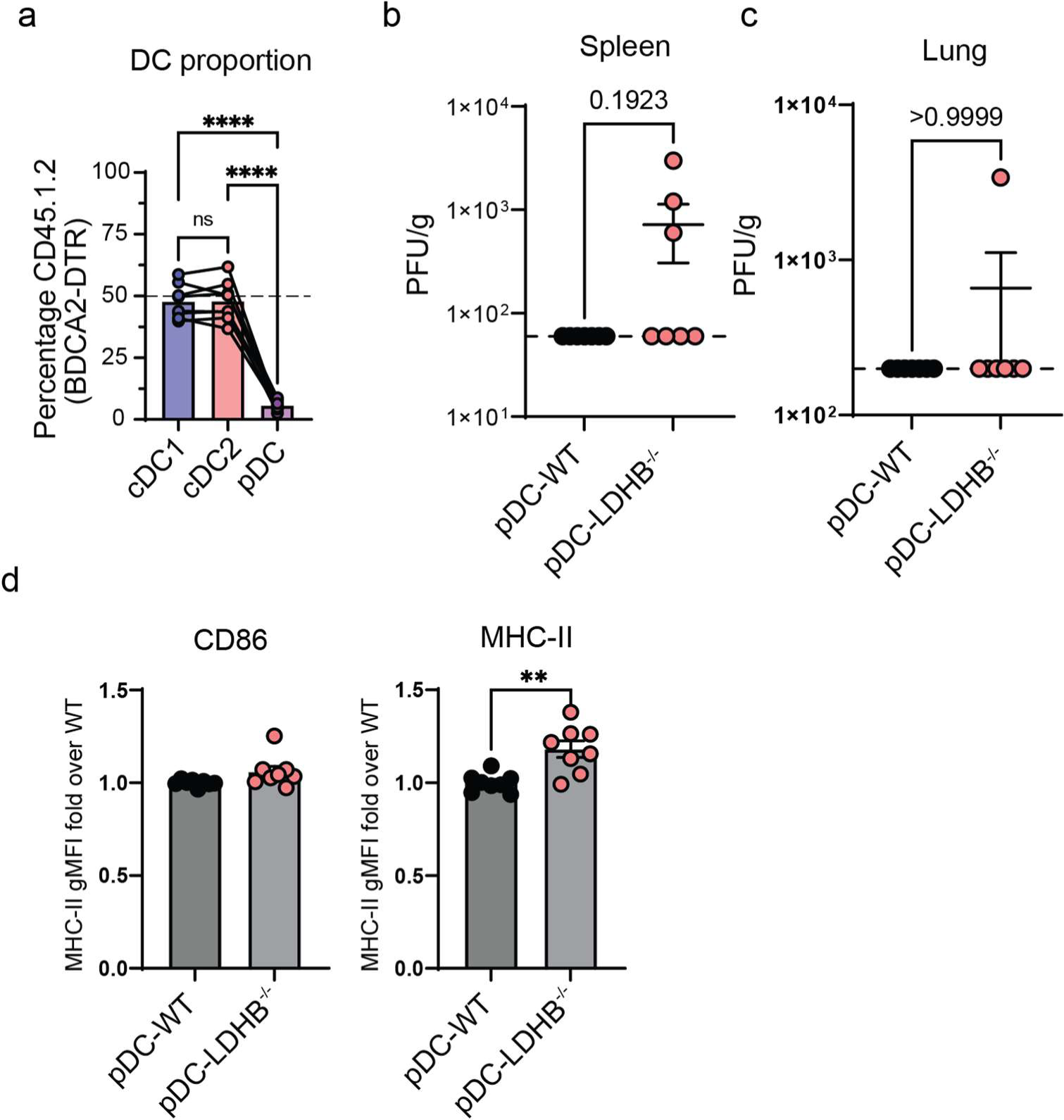
BDCA2-DTR mixed bone marrow chimeras. **(a)** proportion of cDC1 (blue), cDC2 (red), and pDC (purple), from the CD45.1.2 BDCA2-DTR compartment of MHV infected mice described in Fig. 3e after treatment with DT at 48 h.p.i. **(b,c)** Plaque forming units of MHV from the Spleen **(b)** or Lung **(c)** of MHV infected mice described in Fig. 3e at 48 h.p.i. Limit of detection shown as dotted line, samples with no countable plaques are shown at the limit of detection. **(d)** CD86 and MHC-II expression in pDCs isolated from the spleens of the MHV infected mixed BM chimeras as described in Fig. 3e was measured by flow cytometry at 48 h.p.i. Data are pooled from 2-3 independent experiments **(a-d)**. Data are shown as mean ± SEM. Statistics used are One Way ANOVA with Tukey Correction **(a)**. Student’s T Test **(b-d)** p<0.05 *, p<0.01; **, p<0.001 ***, p<0.0001 ****.

**Extended Data Figure 4.**
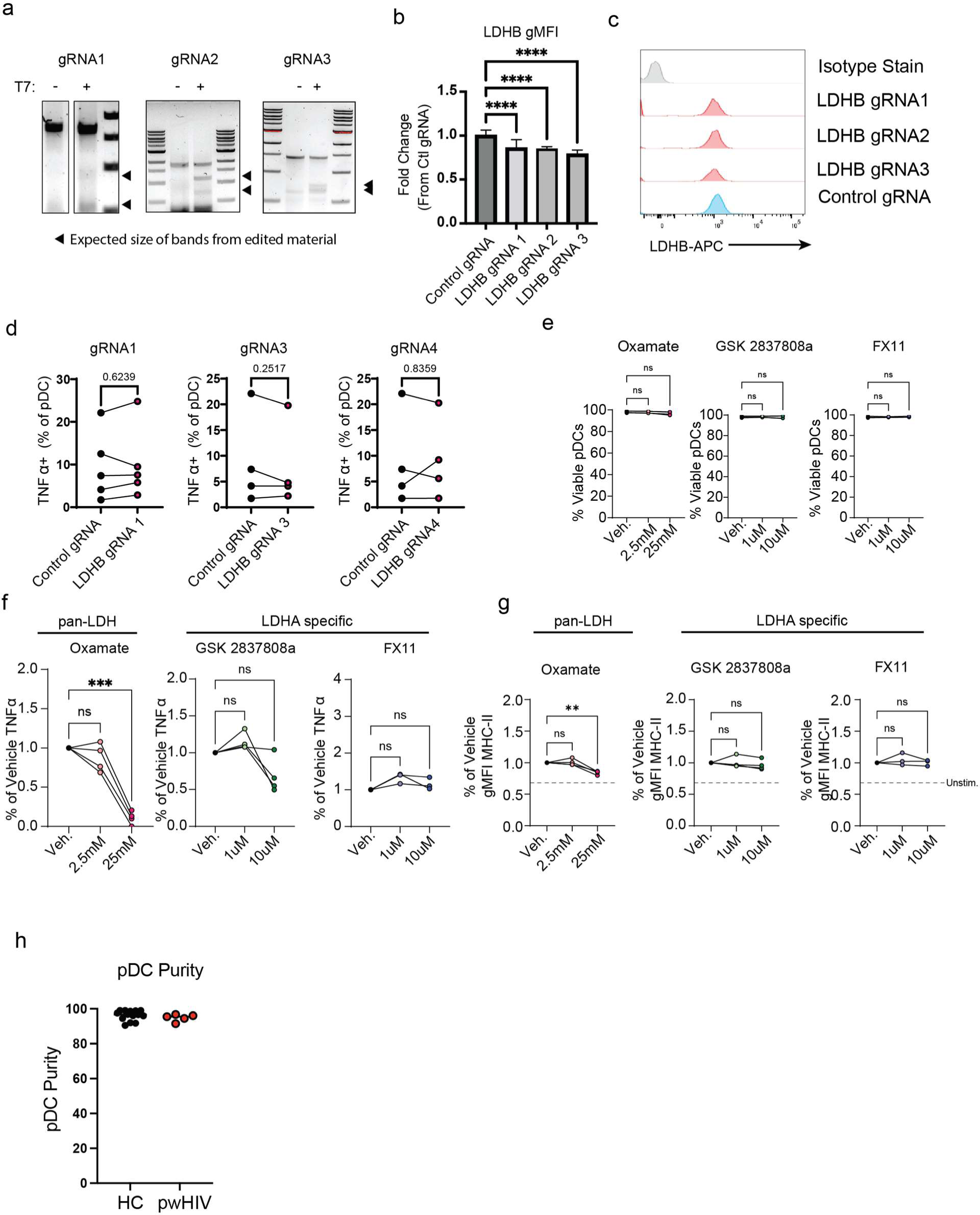
Modulation of LDHB in human pDCs regulates their function. **(a)** Representative gel for T7 assays using DNA isolated from pDC transfected with LDHB targeting gRNA. Arrows represent expected sizes for bands indicating editing. **(b)** gMFI of LDHB in pDCs transfected with LDHB targeting gRNA relative to control gRNA as was measured by flow cytometry. **(c)** Representative histograms from b. **(d)** TNFα production in human pDCs transfected with the indicated Cas9-RNPs after stimulation with R848 as in Fig. 4. **(e)** PBMCs were treated with Oxamate, GSK2837808a, or FX11, stimulated with CpG-A and pDC viability was measured by flow cytometry. **(f,g)** TNFα **(f)** or MHC-II **(g)** expression in pDCs from PBMCs treated with the indicated inhibitors and stimulated with CpG-A for 8 hrs. (**h**) purity of pDC isolated from HC or pwHIV determined by FACS after enrichment. Data are pooled from 3-5 individual donors **(b,d-g)**, representative of 3-5 independent experiments **(a,c)**, or pooled averages from 19 patients quantified across 3 independent experiments **(h)**. Data are shown as mean ± SEM. Statistics used are One Way ANOVA with Tukey Correction **(b,e-g)** or Paired T Test **(d)** p<0.05 *, p<0.01; **, p<0.001 ***, p<0.0001 ****.

**Extended Data Figure 5.**
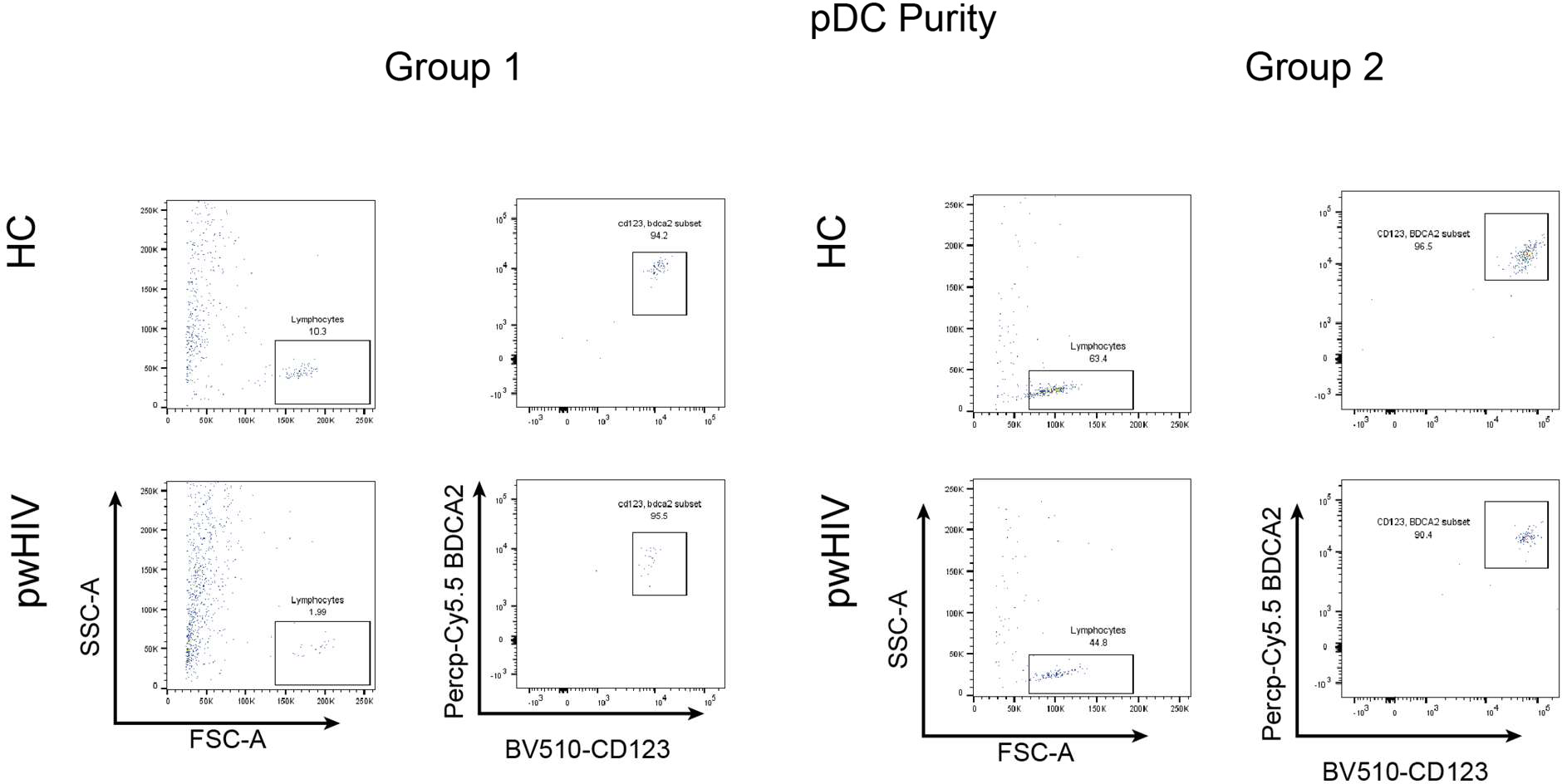
Purification of pDC from pwHIV. Flow cytometry analysis of purified pDC from healthy controls (HC) and people with HIV (pwHIV) for two groups of age/sex matched donors, pDCs are defined as CD123^+^, BDCA2^+^.

**Extended Data Figure 6.**
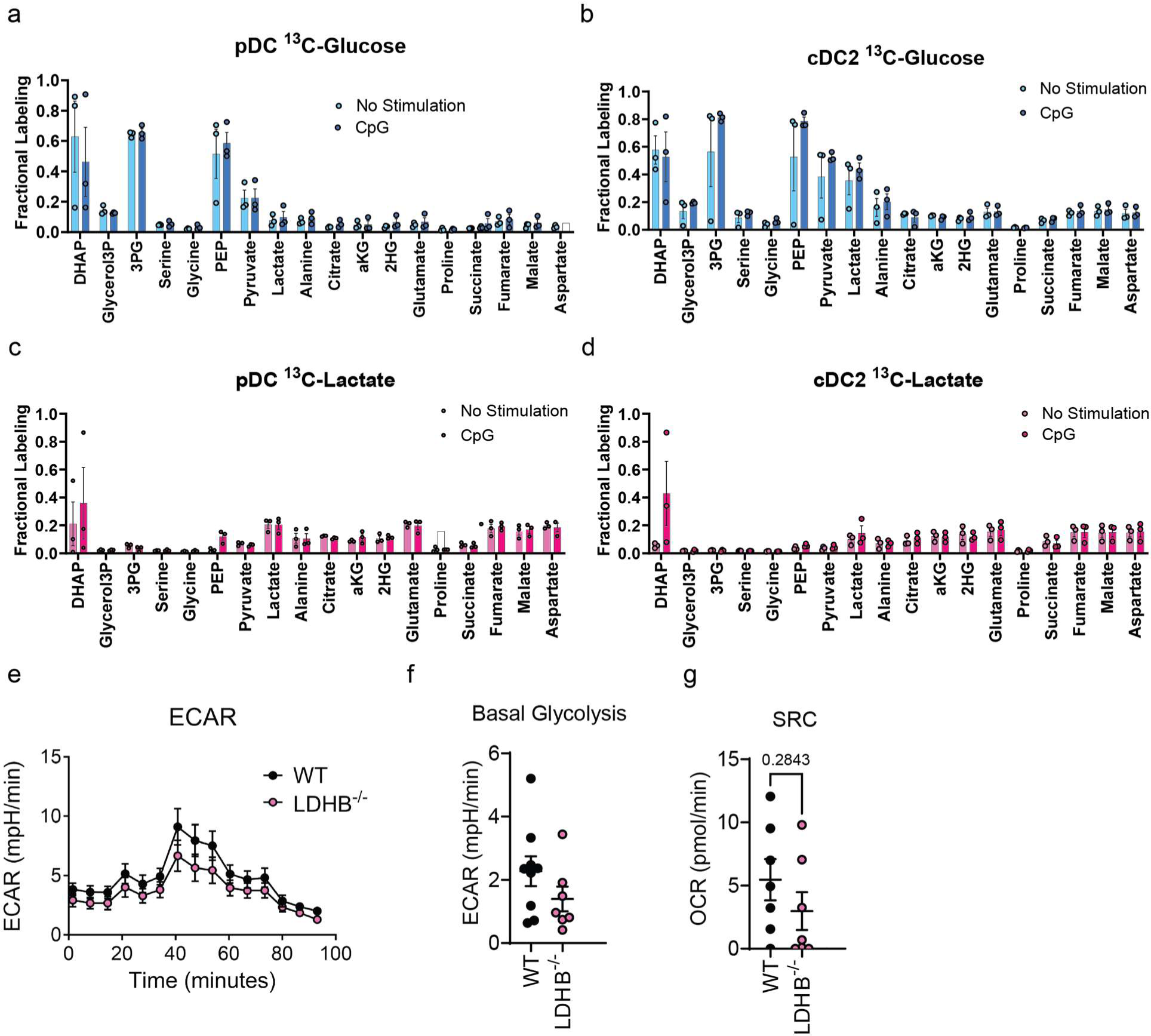
Dendritic Cell Metabolic Profiling. Fractional labeling of glycolysis and TCA cycle intermediates by ^13^C-Glucose **(a,b)** ^13^C-Lactate **(c,d)** in pDCs **(a,c)** and cDC2 **(b,d)** isolated from Flt3L culture. During incubation with ^13^C-sources cells were either unstimulated (light), or stimulated with CpG-A (dark). **(e)** ECAR trace from seahorse of WT (black) and LDHB^-/-^ (pink) pDCs. **(f)** Basal Glycolysis of pDCs from WT (black) and LDHB^-/-^ (pink) mice. **(g)** Spare Respirator Capacity of pDCs from WT (black) and LDHB^-/-^ (pink) mice. Data are pooled from 3-4 independent experiments **(a-g)**. Data are shown as mean ± SEM. Statistics used are Student’s t test **(a-d,f,g)**.

**Extended Data Figure 7.**
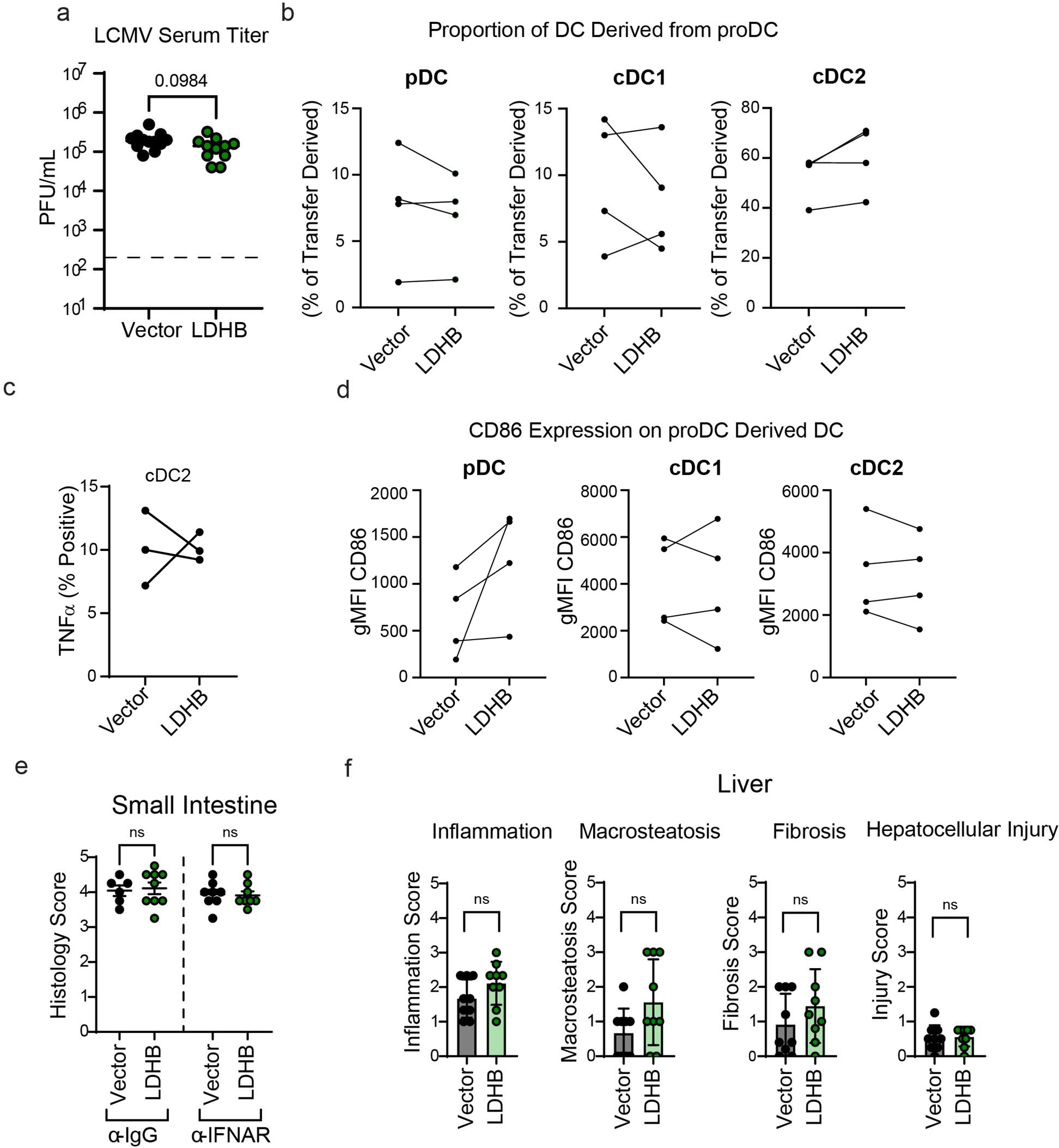
Enforced expression of LDHB in transferred pDCs and infection outcome. **(a)** Viral titers measured by plaque assay in serum day 13.5 p.i. (6 days after transfer of vector or LDHB expressing pro-DCs), as described in Fig. 6b. **(b)** proportion of spleen DCs derived from vector or LDHB expressing pro-DCs after *in vivo* transfer was evaluated by flow cytometry. **(c)** TNFα production in cDC2 derived from vector or LDHB expressing pro-DC transferred into LCMV Cl13 infected animals after CpG-A stimulation for 8 hrs. **(d)** Expression of CD86 on pDC, cDC1, and cDC2 derived from vector or LDHB expressing pro-DCs after stimulation with CpG-A for 8 hrs. **(e,f)** Small intestine **(e)** and Liver **(f)** histology scores after transfer of differentiated pDCs expressing vector or LDHB as described in Fig. 6. Data are pooled from 2-4 experiments **(a-f)**. Statistics used are student’s t test **(a,f)**, paired t test **(b-d)**, and two-way ANOVA with Fisher’s LSD test **(e)**.

## Acknowledgements

We thank Christie Lyn Costanza and Marta Paez-Quinde for help with recruitment of human subjects. This study was supported by NIH grants (AI145314, AI132122, and AI081923) (to E.I.Z.),R50CA252146 (to T.K)), and R50CA283813 (to D.A.S.). The Sanford Burnham Prebys Cancer Metabolism Core is supported in part by Cancer Center support grant P30CA030199. We thank Olga Zagnitko for technical support with GC-MS analysis. We thank Prof. Susan Weiss for providing MHV A59. We also thank Celldex Therapeutics Inc. for providing Flt3L for these studies. We thank applied pathology services (APS) for their work scoring liver injury.

## Contributions

T.T.G. designed, performed, analyzed, and interpreted most experiments, made the figures, and wrote the manuscript. Y.J. performed and analyzed the RNA-seq experiments in combination with microarray data, confirmed TLR7-dependent downregulation of LDHB in exhausted pDCs and identified LDHB as a candidate for pDC exhaustion regulation. M.M. performed and analyzed the microarray experiments in combination with the RNA-Seq data. F.K., K.R.K, C.C., and Z.F. assisted with experiments. Z.F. independently validated IFN-I production in LDHB overexpressing pDCs. Y.F. performed histological scoring. S.S. recruited patients for HIV studies. P.F. provided HIV patient samples, and A.L.C. and E.A. isolated pDCs and performed FACS analysis from HIV patient samples. D.A.S., T.K., and C.M. performed the GC-MS experiments quantifying ^13^C incorporation. Y.J., M.M., Y.F., P.F, D.A.S, and C.M. provided useful feedback on the manuscript. E.I.Z. conceived and supervised the overall study, designed, and interpreted experiments and wrote the manuscript.

## References

1. McNab, F., Mayer-Barber, K., Sher, A., Wack, A. & O’Garra, A. Type I interferons in infectious disease. Nat Rev Immunol 15, 87–103 (2015).

2. Swiecki, M. & Colonna, M. The multifaceted biology of plasmacytoid dendritic cells. Nat Rev Immunol 15, 471–485 (2015).

3. Zuniga, E. I., Liou, L.-Y., Mack, L., Mendoza, M. & Oldstone, M. B. A. Persistent virus infection inhibits type I interferon production by plasmacytoid dendritic cells to facilitate opportunistic infections. Cell Host Microbe 4, 374–86 (2008).

4. Lee, L. N., Burke, S., Montoya, M. & Borrow, P. Multiple Mechanisms Contribute to Impairment of Type 1 Interferon Production during Chronic Lymphocytic Choriomeningitis Virus Infection of Mice. The Journal of Immunology 182, 7178– 7189 (2009).

5. Greene, T. T., Jo, Y. & Zuniga, E. I. Infection and cancer suppress pDC derived IFN-I. Curr Opin Immunol 66, 114–122 (2020).

6. Zuniga, E. I., Macal, M., Lewis, G. M. & Harker, J. A. Innate and Adaptive Immune Regulation During Chronic Viral Infections. Annu Rev Virol. 2, 573–597 (2015).

7. Virgin, H. W., Wherry, E. J. & Ahmed, R. Redefining Chronic Viral Infection. Cell 138, 30–50 (2009).

8. Stelekati, E. & Wherry, E. J. Chronic bystander infections and immunity to unrelated antigens. Cell Host Microbe 12, 458–469 (2012).

9. Attanasio, J. & Wherry, E. J. Costimulatory and Coinhibitory Receptor Pathways in Infectious Disease. Immunity 44, 1052–1068 (2016).

10. Buchbinder, E. I. & Hodi, F. S. Immune-checkpoint blockade — durable cancer control. Nat Rev Clin Oncol 13, 77–78 (2016).

11. Wykes, M. N. & Lewin, S. R. Immune checkpoint blockade in infectious diseases. Nat Rev Immunol 18, 91–104 (2018).

12. Moir, S. & Fauci, A. S. B-cell exhaustion in HIV infection: The role of immune activation. Curr Opin HIV AIDS 9, 472–477 (2014).

13. Zitvogel, L., Galluzzi, L., Kepp, O., Smyth, M. J. & Kroemer, G. Type I interferons in anticancer immunity. Nat Rev Immunol 15, 405–414 (2015).

14. García-Sastre, A. & Biron, C. A. Type 1 interferons and the virus-host relationship: A lesson in détente. Science (1979) 312, 879–882 (2006).

15. Reizis, B. Plasmacytoid Dendritic Cells: Development, Regulation, and Function. Immunity 50, 37–50 (2019).

16. Chiale, C., Greene, T. T. & Zuniga, E. I. Interferon induction, evasion, and paradoxical roles during SARS-CoV-2 infection*. Immunol Rev 309, 12–24 (2022).

17. Fitzgerald-Bocarsly, P. & Feng, D. The role of type I interferon production by dendritic cells in host defense. Biochimie 89, 843–855 (2007).

18. Macal, M. et al. Self-Renewal and Toll-like Receptor Signaling Sustain Exhausted Plasmacytoid Dendritic Cells during Chronic Viral Infection. Immunity 48, 730–744.e5 (2018).

19. Siegal, F. P. et al. Opportunistic infections in acquired immune deficiency syndrome result from synergistic defects of both the natural and adaptive components of cellular immunity. Journal of Clinical Investigation 78, 115–123 (1986).

20. Lopez, C., Fitzgerald, P. A. & Siegal, F. P. Severe Acquired Immune Deficiency Syndrome in Male Homosexuals: Diminished Capacity to Make Interferon-a in Vitro Associated with Severe Opportunistic Infections. Journal of Infectious Diseases 148, 962–966 (1983).

21. Hartmann, E. et al. Identification and functional analysis of tumor-infiltrating plasmacytoid dendritic cells in head and neck cancer. Cancer Res 63, 6478–87 (2003).

22. Sisirak, V. et al. Impaired IFN-α production by plasmacytoid dendritic cells favors regulatory T-cell expansion that may contribute to breast cancer progression. Cancer Res 72, 5188–97 (2012).

23. Labidi-Galy, S. I. et al. Quantitative and functional alterations of plasmacytoid dendritic cells contribute to immune tolerance in ovarian cancer. Cancer Res 71, 5423–34 (2011).

24. Saulep-Easton, D. et al. Cytokine-driven loss of plasmacytoid dendritic cell function in chronic lymphocytic leukemia. Leukemia 28, 2005–2015 (2014).

25. Le Mercier, I. et al. Tumor Promotion by Intratumoral Plasmacytoid Dendritic Cells Is Reversed by TLR7 Ligand Treatment. Cancer Res 73, 4629–4640 (2013).

26. Nierkens, S. et al. Immune adjuvant efficacy of CpG oligonucleotide in cancer treatment is founded specifically upon TLR9 function in plasmacytoid dendritic cells. Cancer Res 71, 6428–37 (2011).

27. Lou, Y. et al. Antitumor Activity Mediated by CpG: The Route of Administration is Critical. Journal of Immunotherapy 34, 279–288 (2011).

28. Tel, J. et al. Natural human plasmacytoid dendritic cells induce antigen-specific T-cell responses in melanoma patients. Cancer Res 73, 1063–75 (2013).

29. Macal, M. et al. Plasmacytoid dendritic cells are productively infected and activated through TLR-7 early after arenavirus infection. Cell Host Microbe 11, 617–30 (2012).

30. Bauer, M. et al. Bacterial CpG-DNA Triggers Activation and Maturation of Human CD11c, CD123 Dendritic Cells 1. The Journal of Immunology 166, 5000–5007 (2001).

31. Bennett, J., Pomaznoy, M., Singhania, A. & Peters, B. A metric for evaluating biological information in gene sets and its application to identify co-expressed gene clusters in PBMC. PLoS Comput Biol 17, e1009459 (2021).

32. Thorndike, R. L. Who belongs in the family? Psychometrika 18, 267–276 (1953).

33. Gu, Z. & Hübschmann, D. H. simplifyEnrichment: A Bioconductor Package for Clustering and Visualizing Functional Enrichment Results. doi:10.1016/j.gpb.2022.04.008.

34. Assil, S. et al. Plasmacytoid Dendritic Cells and Infected Cells Form an Interferogenic Synapse Required for Antiviral Responses. Cell Host Microbe 25, 730–745.e6 (2019).

35. Bruni, D. et al. Viral entry route determines how human plasmacytoid dendritic cells produce type i interferons. Sci Signal 8, ra25 (2015).

36. Décembre, E. et al. Sensing of Immature Particles Produced by Dengue Virus Infected Cells Induces an Antiviral Response by Plasmacytoid Dendritic Cells. PLoS Pathog 10, 1004434 (2014).

37. Feng, Z. et al. Human pDCs preferentially sense enveloped hepatitis A virions. J Clin Invest 125, 169–176 (2015).

38. García-Nicolás, O. et al. Sensing of Porcine Reproductive and Respiratory Syndrome Virus-Infected Macrophages by Plasmacytoid Dendritic Cells. Front Microbiol 7, 771 (2016).

39. Lepelley, A. et al. Innate Sensing of HIV-Infected Cells. PLoS Pathog 7, e1001284 (2011).

40. Wieland, S. F. et al. Human Plasmacytoid Dendritic Cells Sense Lymphocytic Choriomeningitis Virus-Infected Cells In Vitro. J Virol 88, 752–757 (2014).

41. Yun, T. J., et al. Human plasmacytoid dendritic cells mount a distinct antiviral response to virus-infected cells. Sci Immunol 6, (2021).

42. Takahashi, K. et al. Plasmacytoid dendritic cells sense hepatitis C virus-infected cells, produce interferon, and inhibit infection. Proc Natl Acad Sci U S A 107, 7431–7436 (2010).

43. Swiecki, M. et al. Type I interferon negatively controls plasmacytoid dendritic cell numbers in vivo. J Exp Med 208, 2367–2374 (2011).

44. Lee, M. Y. H. et al. Tissue-specific transcriptional profiling of plasmacytoid dendritic cells reveals a hyperactivated state in chronic SIV infection. PLoS Pathog 17, e1009674 (2021).

45. Tu, Z. et al. Cross-linking of CD81 by HCV-E2 protein inhibits human intrahepatic plasmacytoid dendritic cells response to CpG-ODN. Cell Immunol 284, 98–103 (2013).

46. Zhang, S., Kodys, K., Babcock, G. J. & Szabo, G. CD81/CD9 tetraspanins aid plasmacytoid dendritic cells in recognition of hepatitis C virus-infected cells and induction of interferon-alpha. Hepatology 58, 940–949 (2013).

47. Kim, T. W. et al. Transcriptional Repression of IFN Regulatory Factor 7 by MYC Is Critical for Type I IFN Production in Human Plasmacytoid Dendritic Cells. The Journal of Immunology 197, 3348–3359 (2016).

48. Bajwa, G. et al. Cutting Edge: Critical Role of Glycolysis in Human Plasmacytoid Dendritic Cell Antiviral Responses. The Journal of Immunology 196, 2004–2009 (2016).

49. Wu, D. et al. Type 1 Interferons Induce Changes in Core Metabolism that Are Critical for Immune Function. Immunity 44, 1325–1336 (2016).

50. Hurley, H. J. et al. Frontline Science: AMPK regulates metabolic reprogramming necessary for interferon production in human plasmacytoid dendritic cells. J Leukoc Biol 109, 299–308 (2021).

51. Wu, M. et al. Multiparameter metabolic analysis reveals a close link between attenuated mitochondrial bioenergetic function and enhanced glycolysis dependency in human tumor cells. Am J Physiol Cell Physiol 292, 125–136 (2007).

52. Bengsch, B. et al. Bioenergetic Insufficiencies Due to Metabolic Alterations Regulated by the Inhibitory Receptor PD-1 Are an Early Driver of CD8 + T Cell Exhaustion. Immunity 45, 358–373 (2016).

53. Dawson, D. M., Goodfriend, T. L., Kaplan, N. O. & Kaplan, N. O. Lactic Dehydrogenases: Functions of the Two Types. Science (1979) 143, 929–933 (1964).

54. Cahn, R. D., Kaplan, N. O., Levine L. & Zwilling E. Nature and Development of Lactic Dehydrogenases. Science 962–969 https://www.jstor.org/stable/1708677 (1962).

55. Cheng, A. et al. Aurora-A mediated phosphorylation of LDHB promotes glycolysis and tumor progression by relieving the substrate-inhibition effect. Nat Commun 10, (2019).

56. Zdralevic, M. et al. Double genetic disruption of lactate dehydrogenases A and B is required to ablate the ‘Warburg effect’ restricting tumor growth to oxidative metabolism. (2018) doi:10.1074/jbc.RA118.004180.

57. Dickinson, M. E. et al. High-throughput discovery of novel developmental phenotypes. Nature 537, 508–514 (2016).

58. Cervantes-Barragan, L. et al. Plasmacytoid dendritic cells control T-cell response to chronic viral infection. Proceedings of the National Academy of Sciences 109, 3012–3017 (2012).

59. Asselin-Paturel, C. et al. Type I interferon dependence of plasmacytoid dendritic cell activation and migration. J Exp Med 201, 1157 (2005).

60. Swiecki, M., Gilfillan, S., Vermi, W., Wang, Y. & Colonna, M. Plasmacytoid dendritic cell ablation impacts early interferon responses and antiviral NK and CD8(+) T cell accrual. Immunity 33, 955–66 (2010).

61. Channappanavar, R. et al. Dysregulated Type I Interferon and Inflammatory Monocyte-Macrophage Responses Cause Lethal Pneumonia in SARS-CoV-Infected Mice. Cell Host Microbe 19, 181–193 (2016).

62. Greene, T. T. & Zuniga, E. I. Type i interferon induction and exhaustion during viral infection: Plasmacytoid dendritic cells and emerging covid-19 findings. Viruses vol. 13 Preprint at 10.3390/v13091839 (2021).

63. Channappanavar, R. et al. IFN-I response timing relative to virus replication determines MERS coronavirus infection outcomes. Journal of Clinical Investigation 129, 3625–3639 (2019).

64. Cervantes-Barragan, L. et al. Control of coronavirus infection through plasmacytoid dendritic-cell- derived type I interferon. Blood 109, 1131–1137 (2007).

65. Hochrein, H., O’Keeffe, M. & Wagner, H. Human and mouse plasmacytoid dendritic cells. Hum Immunol 63, 1103–1110 (2002).

66. Wong, C. et al. Selective inhibition of the sperm-specific lactate dehydrogenase isozyme-C4 by N-isopropyl oxamate. Biochim Biophys Acta 1343, 16–22 (1997).

67. Brooks, G. A., Dubouchaud, H., Brown, M., Sicurello, J. P. & Eric Butz, C. Role of mitochondrial lactate dehydrogenase and lactate oxidation in the intracellular lactate shuttle. Proc Natl Acad Sci U S A 96, 1129–1134 (1999).

68. Billiard, J. et al. Quinoline 3-sulfonamides inhibit lactate dehydrogenase A and reverse aerobic glycolysis in cancer cells. (2013) doi:10.1186/2049-3002-1-19.

69. Yu, Y., et al. Selective active site inhibitors of human lactate dehydrogenases A 4, B 4, and C 4. (2001).

70. Annas, D. et al. Synthesis and initial screening of lactate dehydrogenase inhibitor activity of 1,3-benzodioxole derivatives. Scientific Reports 2020 10:1 10, 1–9 (2020).

71. Mohammad, G. H. et al. Targeting Pyruvate Kinase M2 and Lactate Dehydrogenase A Is an Effective Combination Strategy for the Treatment of Pancreatic Cancer. Cancers (Basel) 11, (2019).

72. Mediani, L. et al. Reversal of the glycolytic phenotype of primary effusion lymphoma cells by combined targeting of cellular metabolism and PI3K/Akt/ mTOR signaling. Oncotarget 7,.

73. Feldman, S. et al. Decreased Interferon-α Production in HIV-Infected Patients Correlates with Numerical and Functional Deficiencies in Circulating Type 2 Dendritic Cell Precursors. Clinical Immunology 101, 201–210 (2001).

74. Zhang, Z. et al. Differential Restoration of Myeloid and Plasmacytoid Dendritic Cells in HIV-1-Infected Children after Treatment with Highly Active Antiretroviral Therapy. The Journal of Immunology 176, 5644–5651 (2006).

75. Tilton, J. C. et al. Human Immunodeficiency Virus Viremia Induces Plasmacytoid Dendritic Cell Activation In Vivo and Diminished Alpha Interferon Production In Vitro. J Virol 82, 3997–4006 (2008).

76. Schwartz, J. A. et al. Tim-3 is a Marker of Plasmacytoid Dendritic Cell Dysfunction during HIV Infection and Is Associated with the Recruitment of IRF7 and p85 into Lysosomes and with the Submembrane Displacement of TLR9. The Journal of Immunology 198, 3181–3194 (2017).

77. Goutagny, N. et al. Quantification and Functional Analysis of Plasmacytoid Dendritic Cells in Patients with Chronic Hepatitis C Virus Infection. J Infect Dis 189, 1646–1655 (2004).

78. Kanto, T. et al. Reduced Numbers and Impaired Ability of Myeloid and Plasmacytoid Dendritic Cells to Polarize T Helper Cells in Chronic Hepatitis C Virus Infection. J Infect Dis 190, 1919–1926 (2004).

79. Ulsenheimer, A. et al. Plasmacytoid dendritic cells in acute and chronic hepatitis C virus infection. Hepatology 41, 643–651 (2005).

80. Lai, W. K. et al. Hepatitis C is associated with perturbation of intrahepatic myeloid and plasmacytoid dendritic cell function. J Hepatol 47, 338–347 (2007).

81. Dolganiuc, A. et al. Hepatitis C Virus (HCV) Core Protein-Induced, Monocyte- Mediated Mechanisms of Reduced IFN-α and Plasmacytoid Dendritic Cell Loss in Chronic HCV Infection. The Journal of Immunology 177, 6758–6768 (2006).

82. van der Molen, R. G. et al. Functional impairment of myeloid and plasmacytoid dendritic cells of patients with chronic hepatitis B. Hepatology 40, 738–746 (2004).

83. Duan, X.-Z. et al. Decreased Frequency and Function of Circulating Plasmocytoid Dendritic Cells (pDC) in Hepatitis B Virus Infected Humans. J Clin Immunol 24, 637–646 (2004).

84. Xie, Q. et al. Patients with chronic hepatitis B infection display deficiency of plasmacytoid dendritic cells with reduced expression of TLR9. Microbes Infect 11, 515–523 (2009).

85. Soumelis, V. et al. Depletion of circulating natural type 1 interferon-producing cells in HIV-infected AIDS patients. Blood 98, 906–912 (2001).

86. Chehimi, J. et al. Persistent Decreases in Blood Plasmacytoid Dendritic Cell Number and Function Despite Effective Highly Active Antiretroviral Therapy and Increased Blood Myeloid Dendritic Cells in HIV-Infected Individuals. The Journal of Immunology 168, 4796–4801 (2002).

87. Meyers, J. H. et al. Impact of HIV on Cell Survival and Antiviral Activity of Plasmacytoid Dendritic Cells. PLoS One 2, e458 (2007).

88. Naik, S. H. et al. Development of plasmacytoid and conventional dendritic cell subtypes from single precursor cells derived in vitro and in vivo. (2007) doi:10.1038/ni1522.

89. Malleret, B. et al. Primary infection with simian immunodeficiency virus: plasmacytoid dendritic cell homing to lymph nodes, type I interferon, and immune suppression. Blood 112, 4598–4608 (2008).

90. Bruel, T. et al. Plasmacytoid Dendritic Cell Dynamics Tune Interferon-Alfa Production in SIV-Infected Cynomolgus Macaques. PLoS Pathog 10, e1003915 (2014).

91. Szabo, G. & Dolganiuc, A. Subversion of plasmacytoid and myeloid dendritic cell functions in chronic HCV infection. Immunobiology 210, 237–247 (2005).

92. Arunachalam, P. S. et al. Systems biological assessment of immunity to mild versus severe COVID-19 infection in humans. Science (1979) 369, 1210–1220 (2020).

93. Barber, D. L. et al. Restoring function in exhausted CD8 T cells during chronic viral infection. Nature 439, 682–687 (2006).

94. Macleod, B. L. et al. A network of immune and microbial modifications underlies viral persistence in the gastrointestinal tract. Journal of Experimental Medicine 217, (2020).

95. Bunin, A. et al. Protein Tyrosine Phosphatase PTPRS Is an Inhibitory Receptor on Human and Murine Plasmacytoid Dendritic Cells. Immunity 43, 277–288 (2015).

96. Wang, Y. et al. Timing and Magnitude of Type I Interferon Responses by Distinct Sensors Impact CD8 T Cell Exhaustion and Chronic Viral Infection. Cell Host Microbe 11, 631–642 (2012).

97. Sheehan, K. C. F. et al. Blocking monoclonal antibodies specific for mouse IFN-α/β receptor subunit 1 (IFNAR-1) from mice immunized by in vivo hydrodynamic transfection. Journal of Interferon and Cytokine Research 26, 804–819 (2006).

98. Chauvin, J.-M. & Zarour, H. M. Open access TIGIT in cancer immunotherapy. J Immunother Cancer 8, 957 (2020).

99. Ghosh, H. S., Cisse, B., Bunin, A., Lewis, K. L. & Reizis, B. Continuous Expression of the Transcription Factor E2-2 Maintains the Cell Fate of Mature Plasmacytoid Dendritic Cells. Immunity 33, 905–916 (2010).

100. Cisse, B. et al. Transcription factor E2-2 is an essential and specific regulator of plasmacytoid dendritic cell development. Cell 135, 37–48 (2008).

101. Cao, W. et al. Toll-like receptor-mediated induction of type I interferon in plasmacytoid dendritic cells requires the rapamycin-sensitive PI(3)K-mTOR- p70S6K pathway. Nat Immunol 9, 1157–1164 (2008).

102. Katarzyna Grzes, A. M., et al. Plasmacytoid dendritic cell activation is dependent on coordinated expression of distinct amino acid transporters. (2021) doi:10.1016/j.immuni.2021.10.009.

103. Donovan, M. L., Schultz, T. E., Duke, T. J. & Blumenthal, A. Type I interferons in the pathogenesis of tuberculosis: Molecular drivers and immunological consequences. Front Immunol 8, 1633 (2017).

104. Parker, D., Planet, P. J., Soong, G., Narechania, A. & Prince, A. Induction of Type I Interferon Signaling Determines the Relative Pathogenicity of Staphylococcus aureus Strains. PLoS Pathog 10, e1003951 (2014).

105. Auerbuch, V., Brockstedt, D. G., Meyer-Morse, N., O’Riordan, M. & Portnoy, D. A. Mice Lacking the Type I Interferon Receptor Are Resistant to Listeria monocytogenes. Journal of Experimental Medicine 200, 527–533 (2004).

106. Boxx, G. M. & Cheng, G. The Roles of Type i Interferon in Bacterial Infection. Cell Host Microbe 19, 760–769 (2016).

107. García-Sastre, A. Ten Strategies of Interferon Evasion by Viruses. Cell Host Microbe 22, 176–184 (2017).

108. Wilson, E. B. et al. Blockade of chronic type I interferon signaling to control persistent LCMV infection. Science (1979) 340, 202–207 (2013).

109. Teijaro, J. R. et al. Persistent LCMV Infection Is Controlled by Blockade of Type I Interferon Signaling. Science (1979) 340, 207–211 (2013).

110. Psarras, A. et al. Functionally impaired plasmacytoid dendritic cells and non- haematopoietic sources of type I interferon characterize human autoimmunity. Nature Communications 2020 11:1 11, 1–18 (2020).

111. Ahmed, R. Selection of genetic variants of lymphocytic choriomeningitis virus in spleens of persistently infected mice. Role in suppression of cytotoxic T lymphocyte response and viral persistence. Journal of Experimental Medicine 160, 521–540 (1984).

112. Hirano, N., Fujiwara, K. & Matumoto, M. Mouse Hepatitis Virus (MHV-2): Plaque Assay and Propagation in Mouse Cell Line DBT Cells. Jpn J Microbiol 20, 219–225 (1976).

113. Gilliet, M. et al. The Development of Murine Plasmacytoid Dendritic Cell Precursors Is Differentially Regulated by FLT3-ligand and Granulocyte/Macrophage Colony-Stimulating Factor. J Exp Med 195, 953–958 (2002).

114. Scott, D. A. Analysis of Melanoma Cell Glutamine Metabolism by Stable Isotope Tracing and Gas Chromatography-Mass Spectrometry. in 91–110 (2021). doi:10.1007/978-1-0716-1205-7_7.

115. Labarta-Bajo, L. et al. Type I IFNs and CD8 T cells increase intestinal barrier permeability after chronic viral infection. (2020) doi:10.1084/jem.20192276.

116. Erben, U. et al. A guide to histomorphological evaluation of intestinal inflammation in mouse models. Int J Clin Exp Pathol 7, 4557 (2014).

117. Kleiner, D. E. et al. Design and validation of a histological scoring system for nonalcoholic fatty liver disease. Hepatology 41, 1313–1321 (2005).

